# Modest functional diversity decline and pronounced composition shifts of microbial communities in a uranium-contaminated aquifer

**DOI:** 10.1101/2024.12.02.626466

**Authors:** Yupeng Fan, Dongyu Wang, Joy X. Yang, Daliang Ning, Zhili He, Ping Zhang, Andrea M. Rocha, Naijia Xiao, Jonathan P. Michael, Katie F. Walker, Dominique C. Joyner, Chongle Pan, Michael W. W. Adams, Matthew W. Fields, Eric J. Alm, David A. Stahl, Terry C. Hazen, Paul D. Adams, Adam P. Arkin, Jizhong Zhou

## Abstract

**Background:** Microbial taxonomic diversity declines with increased environmental stress. Yet, few studies have explored whether phylogenetic and functional diversities track taxonomic diversity along the stress gradient. Here, we investigated bacterial communities within an aquifer in Oak Ridge, Tennessee, USA, which is characterized by a broad spectrum of stressors, including extremely high levels of nitrate, heavy metals like cadmium and chromium, radionuclides such as uranium, and extremely low pH (<3).

**Results:** Both taxonomic and phylogenetic α-diversities were reduced in the most impacted wells, while the decline in functional α-diversity was modest and statistically insignificant, indicating a more robust buffering capacity to environmental stress. Differences in functional gene composition (i.e., functional β-diversity) were pronounced in highly contaminated wells, while convergent functional gene composition was observed in uncontaminated wells. The relative abundances of most carbon degradation genes were decreased in contaminated wells, but genes associated with denitrification, adenylylsulfate reduction, and sulfite reduction were increased. Compared to taxonomic and phylogenetic compositions, environmental variables played a more significant role in shaping functional gene composition, suggesting that niche selection could be more closely related to microbial functionality than taxonomy.

**Conclusions:** Overall, we demonstrated that despite a reduced taxonomic α-diversity, microbial communities under stress maintained functionality underpinned by environmental selection.

## INTRODUCTION

Microorganisms are adversely affected by environmental stressors such as pH (Power et al., 2018), salinity (Ruhl et al., 2018), aridity (Maestre et al., 2015), temperature (Power et al., 2018), antibiotics (Gao et al., 2020) and heavy metals (Chen et al., 2014), leading to common observations that the number of microbial species declines under extreme conditions. Most of previous research, however, has predominantly focused on microbial taxonomy, neglecting to comprehensively assess the entire functional potentials of microbial communities. Consequently, it remains elusive whether microbial functional α-diversity mirrors the patterns observed in taxonomic α-diversity across various environmental stressors. A positive correlation between these two measures is often assumed, implying that functional α-diversity may increase linearly with taxonomic α-diversity. This assumption received partial support from a study on the eastern Tibetan Plateau, where aridity stress concurrently reduced both functional and taxonomic α-diversities, albeit with a weak correlation (Song et al., 2019). Alternatively, functional α-diversity may increase rapidly with low α-taxonomic diversity but saturate with high taxonomic α-diversity, showing a non-linear relationship (Nielsen et al., 2011).

While microbial α-diversity is measured by the number of taxa and their abundance within a community, microbial β-diversity is defined as the variation in community composition between two communities, often expressed through pairwise dissimilarity (Whittaker, 1972). Strong, positive linear correlations between taxonomic and functional gene β-diversity of microbial communities were observed in soil (Fierer et al., 2013) and marine ecosystems (Galand et al., 2018). Conversely, environmental conditions strongly affected the functional gene compositions of global marine microbial communities but only weakly affected taxonomic composition, indicating a decoupling of functionality from taxonomy (Louca et al., 2016; Ma et al., 2019). The lack of correlation between microbial functionality and taxonomy was also observed in the soil mycobiome of the North American continent (Talbot et al., 2014).

Despite significant advancements in environmental microbiome research, there remains a notable gap in generalizable insights into how microbial α- and β-diversities, particularly α- and β-functional diversities, react to various stressors (Rocca et al., 2018). The Anna Karenina Principle, which suggests that disease-associated microbial communities in hosts under stress of disease are more dissimilar than those of healthy ones, has recently been proposed as a framework for understanding microbial dynamics within the animal (Zaneveld et al., 2017) or plant hosts (Arnault et al., 2023). However, It remains an open question whether the principle holds in aquifer microbial communities under stress.

While high levels of heavy metals restrict certain functional properties of bacterial communities (Zhou et al., 2014), few research has yet quantitatively assessed functional diversity across a broad spectrum of heavy metal concentrations, where dramatic changes in species diversity are evident. We, therefore, selected a range of aquifer samples spanning from 0 to 17 mg/L in uranium concentrations, 0 to 9,000 mg/L in nitrate concentrations, and 3.4 to 7.3 in pH. These samples were collected from a legacy waste site with deposition of nitric acid-solubilized uranium waste between 1951 and 1983, along with mixed metal and organic wastes from other facilities of the US Department of Energy. We analyzed bacterial taxonomic and phylogenetic diversities via 16S rRNA gene amplicon sequencing, and functional diversity or functional gene diversity via metagenome shotgun sequencing. We aimed to test the hypothesis that functional diversity mirrors taxonomic or phylogenetic diversities in response to environmental stressors. Specifically, we investigated whether microbial communities in contaminated wells exhibit distinct characteristics compared to those in uncontaminated wells, providing a testbed of the Anna Karenina Principle in aquifer microbial communities under stress.

## RESULTS

### Environmental variables

The levels of conductivity, dissolved nitrous oxide (N_2_O), chloride (Cl^−^), manganese (Mn), and cadmium (Cd) were higher in high-contaminated wells compared to other wells, while the pH levels were lower (*p* < 0.05, Table S1). Additionally, there were higher concentrations of dissolved organic carbon, dissolved carbon dioxide (CO_2_), nitrate (NO_3_ ^−^), sulfate (SO ^2-^), ferrous, potassium (K), calcium (Ca), barium (Ba), aluminum (Al), silver (Ag), iron(Fe), zinc (Zn), strontium (Sr), and uranium (U) along with lower dissolved nitrogen concentrations, dissolved oxygen concentrations, and dissolved methane concentrations in high-contaminated wells than in other wells, though some of these differences were statistically insignificant (*p* > 0.05). The dispersion of environmental variables significantly increased with increased contamination (Fig. S2).

Nitrite (NO ^−^) concentrations in the supernatant, sodium (Na), and magnesium (Mg) concentrations were higher in high-contaminated wells than in mid-contaminated wells, but not detectable in other wells. Nitrate (NO ^−^) concentrations in the supernatant, cobalt (Co), chromium (Cr), gallium (Ga), lithium (Li), and nickel (Ni) were higher in high-contaminated wells compared to mid- and low-contaminated wells, but not detectable in uncontaminated wells. Higher concentrations of arsenic (As), beryllium (Be), cesium (Cs), copper (Cu), and lead (Pb) were found in high-contaminated wells compared to low-contaminated wells, but not detected in other wells.

### Microbial α-, β- and γ-diversities

The taxonomic, phylogenetic, and functional α-diversities were the lowest in high-contaminated wells, while they were the highest in mid-contaminated wells (Fig. 1). When compared to uncontaminated wells, the taxonomic α-diversities in high-contaminated wells were reduced by 85% (*p* = 0.025), and the phylogenetic α-diversities were reduced by 81% (*p* = 0.018, Figure 1). In contrast, functional α-diversities were not significantly different between high-contaminated wells and uncontaminated wells, with a smaller decrease of 55% on average.

**Fig. 1.**
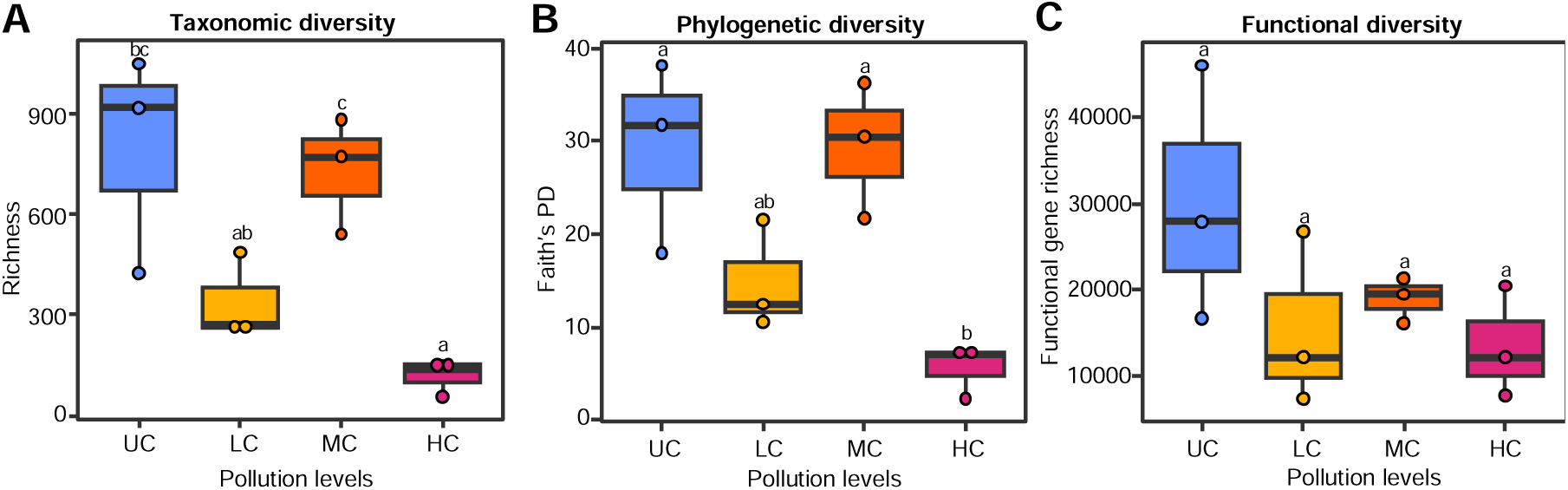
Microbial α-diversity of aquifer samples in uncontaminated wells (UC), low-contaminated wells (LC), mid-contaminated wells (MC), and high-contaminated wells (HC) calculated by richness for (A) taxonomic diversity, (B) phylogenetic diversity, and (C) functional diversity. Letters represent the difference between groups which was determined by ANOVA (significant difference level: p < 0.05).

The taxonomic, phylogenetic, and functional compositions of microbial communities were well separated among uncontaminated, low-, mid-, and high-contaminated wells, as indicated by Non-metric Multidimensional Scaling (NMDS, Fig. 2A-C). To further explore these differences, three permutational tests of dissimilarity (Adonis, MRPP, and ANOSIM) were conducted, which revealed significant differences among the four groups of wells (*p* < 0.001, Table 1). Interestingly, the microbial taxonomic and phylogenetic compositions in high-contaminated wells had lower dispersion compared to the uncontaminated and low-contaminated wells (though not statistically significant, *p* > 0.1 by the permutational dispersion test, Fig. 2D), suggesting that they were more similar in high-contaminated wells. Conversely, microbial functional compositions in high-contaminated wells displayed the highest community dispersion values, indicating a pattern of microbial functional heterogeneity induced in high-contaminated wells (*p* = 0.013 by the permutational dispersion test, Fig. 2D).

**Fig. 2.**
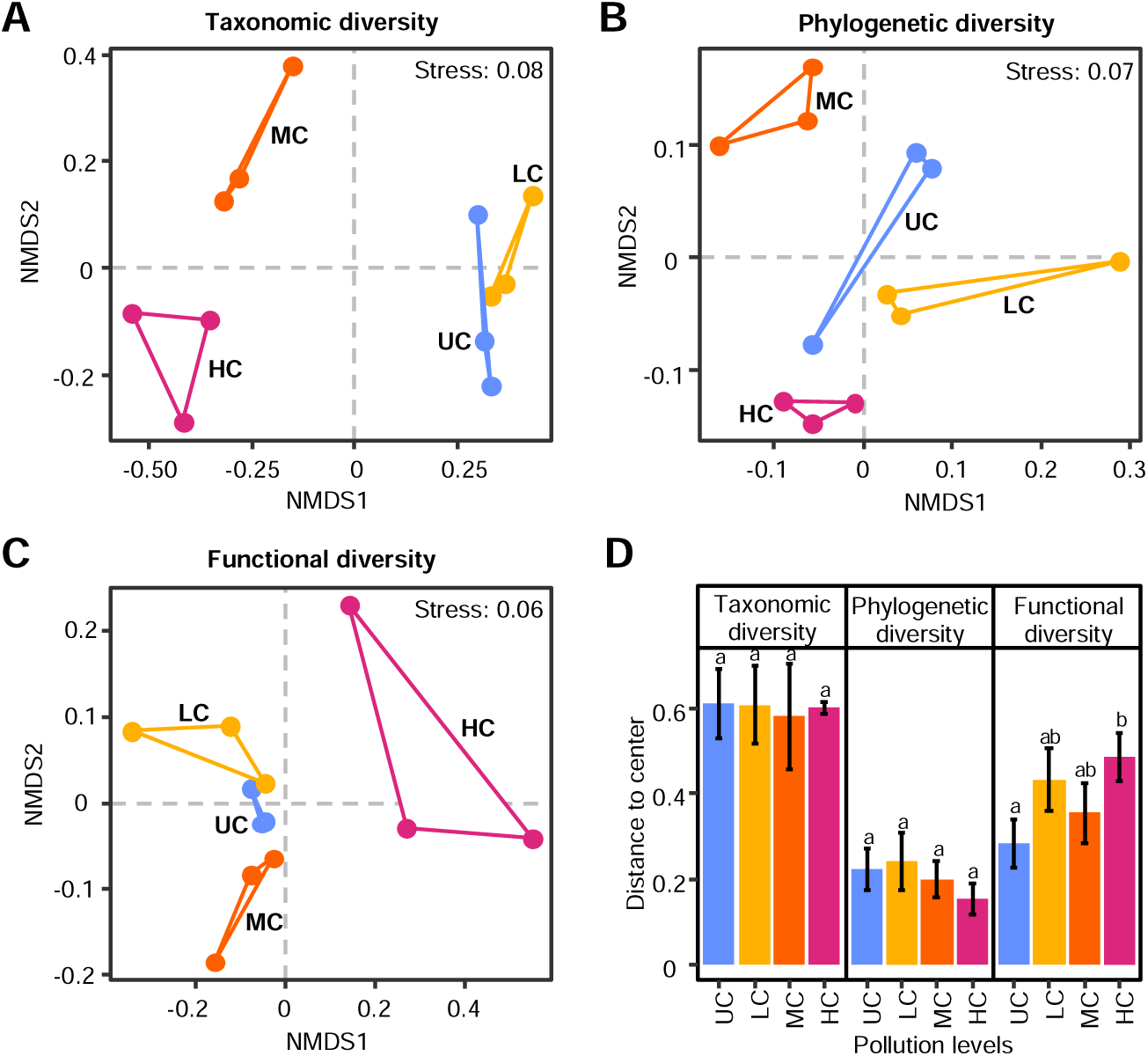
Microbial β-diversity of aquifer microbial communities for different diversity indices in uncontaminated wells (UC), low-contaminated wells (LC), mid-contaminated wells (MC), and high-contaminated wells (HC). Non-metric multidimensional scaling (NMDS) plots based on weighted Bray-Curties index for (A) taxonomic and (C) functional diversities, normalized weighted Unifrac (phylogenetic Bray-Curties) for phylogenetic diversity (B). Dispersion test (D) based on weighted Bray-Curties index for taxonomic and functional diversities, normalized weighted Unifrac (phylogenetic Bray-Curties) for phylogenetic diversity.

**Table 1.** Significance tests of the groundwater microbial communities.

|  | Adonis |  | ANOSIM |  | MRPP |  |
| --- | --- | --- | --- | --- | --- | --- |
| | *F* | *p* | *R* | *p* | $\delta$ | *p* |
| Taxonomic diversity | 1.91 | 0.001 | 0.92 | 0.001 | 0.85 | 0.001 |
| Phylogenetic diversity | 2.43 | 0.001 | 0.57 | 0.001 | 0.33 | 0.001 |
| Functional diversity | 2.19 | 0.001 | 0.46 | 0.001 | 0.57 | 0.002 |

When analyzing γ-diversities, the taxonomic, phylogenetic, and functional γ-diversities in high-contaminated wells were lower than uncontaminted wells, which were similar with those of α-diversities. However, a notable exception was found in the phylogenetic γ-diversity of mid-contaminated wells, which was higher than that in uncontaminated wells (Fig. S3).

### Bacterial taxa

Proteobacteria was the most abundant phylum in high-contaminated wells, accounting for 74% of the relative abundance (Table S2A). In comparison, the average relative abundance of Proteobacteria in other wells was only 21%. Bacterial candidate phylum WPS-2, also known as Eremiobacterota, was higher in high-contaminated wells than others, accounting for 12% of the relative abundance. Acidobacteria was also abundant in mid- and high-contaminated wells, accounting for 5-6% of the relative abundance.

*Rhodanobacter*, a Proteobacteria genus well known for denitrification (Green et al., 2012), was the most abundant in high-contaminated wells (Table S2B), reaching an abundance of 80% in the FW106 well. In comparison, the relative abundance of *Rhodanobacter* in other wells was less than 1%. The second most abundant genus in high-contaminated wells belonged to *Candidatus* phylum Eremiobacterota. The third most abundant genus in high-contaminated wells was *Sulfurifustis*, a genus of sulfur-oxidizing bacteria affiliated with γ-Proteobacteria (Kojima et al., 2015), accounting for 9% of the relative abundance. Other genera of sulfur-oxidizing bacteria, including *Sulfuricurvum* and *Sulfuritalea*, were detected in all wells except for high-contaminated ones.

The genera of nitrifying bacteria and archaea were abundant in certain wells. *Nitrosarchaeum*, a genus of ammonia-oxidizing archaeon, comprised 19% of relative abundance in low-contaminated wells but was only 0.032% to 0.085% in uncontaminated and 0% to 0.364% in mid-contaminated wells. The ammonia-oxidizing bacteria GOUTA6 was detected in all wells, with a relative abundance of 5% in mid-contaminated wells. The nitrite-oxidizing bacteria (NOB) genus *Nitrospira* was present in all wells except high-contaminated wells and accounted for 5% of the relative abundance in low-contaminated wells. The α-Proteobacteria genus *Reyranella*, which produces acetic acid during respiration, was present in all wells, with the highest relative abundance (1.06% to 14.45%) in low-contaminated wells. The methane oxidation bacteria genus *Candidatus Methylomirabilis* was detected in contaminated wells, with the highest relative abundance (0.28% to 14.32%) in mid-contaminated wells.

### Functional gene categories

We used a shotgun metagenomic assembly approach to profile the functional genes. The assembled contigs were annotated by the pathway maps of the KEGG database, in which most annotated pathways in contaminated wells were not significantly different from uncontaminated wells (Fig. S4). Therefore, we used EcoFun-MAP to further annotate the shotgun metagenomic data.

### Carbon degradation genes

We carried out response ratio analyses to reveal carbon degradation genes that were statistically different in relative abundances among uncontaminated, low-, mid-, and high-contaminated wells (*p* < 0.05, Fig. 3A). Most carbon degradation genes were decreased in contaminated wells, including *amyA* encoding α-amylase that hydrolyzes starch and glycogen, *xylA* encoding xylose isomerase that hydrolyzes hemicellulose, endochitinase and exochitinase genes that degrades chitin, *rgaE* encoding acetylesterase that hydrolyzes pectin, *vanA* encoding vanillate monooxygenase that degrades vanillin and lignin, *vdh* encoding vanillin dehydrogenase that degrades vanillin and lignin, and phenol oxidase gene that hydrolyzes lignin. Among them, *ara* encoding l-arabinofuranosidase that degrades hemicellulose decreased with increased contamination, suggesting that its relative abundance was sensitive to environmental contamination.

**Fig. 3.**
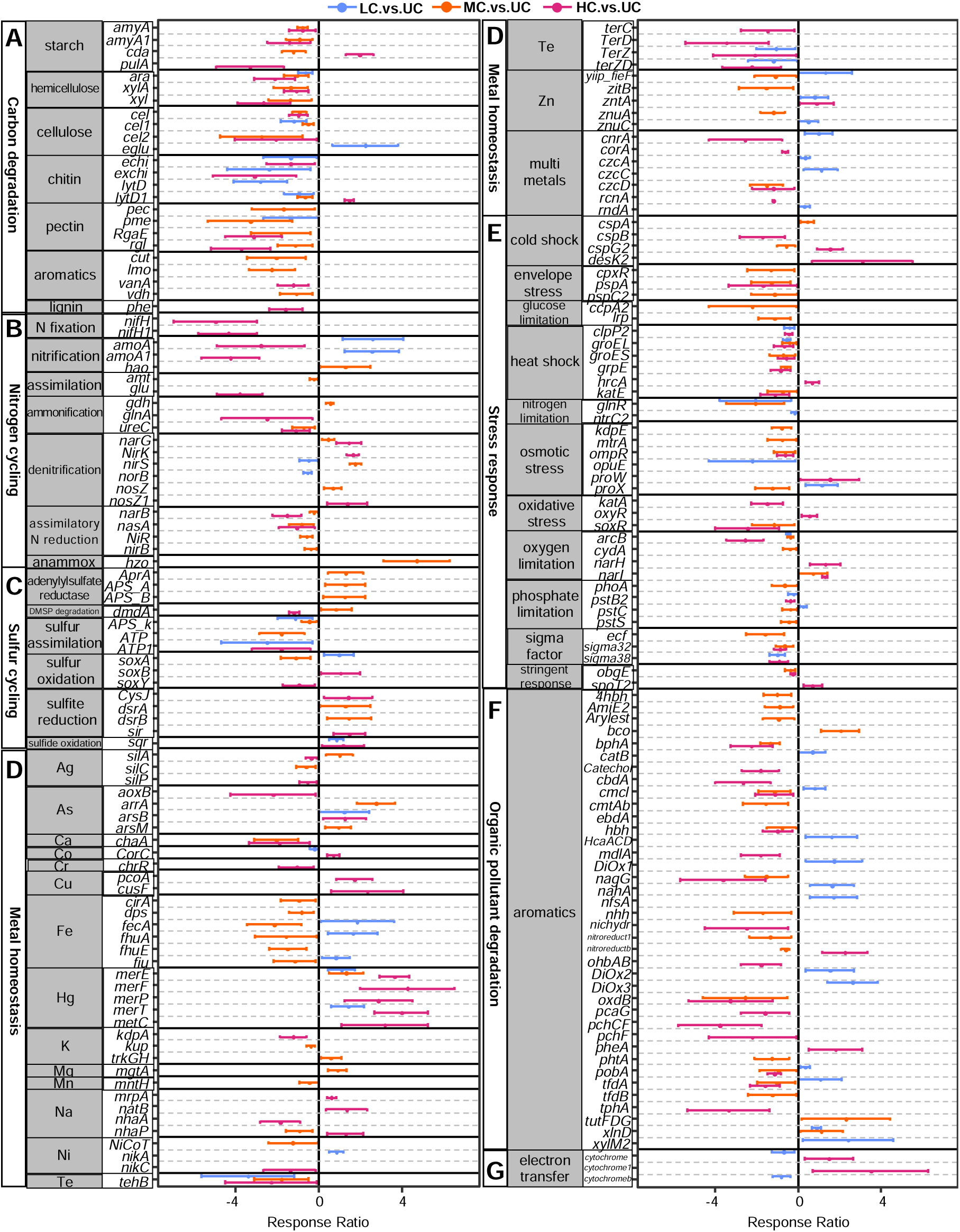
Differences in relative gene abundance for 12 aquifer samples functional community based on response ratio. (A) Carbon degradation genes, (B) Nitrogen cycling genes, (C) Sulfur cycling genes, (D) metal homeostatic genes, (E) stress response genes, (F) organic pollutant degradation genes, and (G) electron transfer genes. All genes presented here are significantly different from those in unpolluted wells, judged by a 95% confidence interval.

### Nitrogen cycling genes

The relative abundances of most denitrification genes, including *narG* encoding nitrate reductase, *nirK* and *nirS* encoding nitrite reductase, and *nosZ* encoding nitrous oxide reductase were increased in mid- and high-contaminated wells (Fig. 3B), which corresponded with high nitrate concentrations in those wells (Table S1). In contrast, biomarker genes of nitrogen fixation (*nifH* encoding the subunit of the Fe protein (Kp2) component of nitrogenase) and nitrification (*amoA* encoding ammonia monooxygenase subunit A) were decreased in high-contaminated wells (Fig. 3B), suggesting that functional potentials of nitrogen fixation and nitrification were reduced by high contamination.

### Sulfur cycling genes

Sulfite reduction genes, including *cysJ, dsrA, dsrB,* and *sir* encoding various sulfite reductase, were increased in mid- and high-contaminated wells (Fig. 3C), consistent with more anaerobic environments in those wells (Table S1). However, sulfur assimilation genes, including APS kinase, ATP sulfurylase in protists, and ATP sulfurylase, were decreased (Fig. 3C), which may suggest a stress response to maintain energy metabolism at the expense of growth.

### Metal homeostasis genes

Many metal homeostasis genes were increased in contaminated wells (Figure 3D), including *merE*, *merF, merP, merT*, and *metC* encoding mercury transporter; *fiu, fhuA*, and *fecA* encoding TonB-dependent receptor that mediates substrate-specific transport across the outer membrane; *arrA* encoding arsenate respiratory reductase*, arsB* encoding an arsenical pump membrane protein, and *arsM* encoding arsenite S-adenosylmethyltransferase; *pcoA* encoding copper resistance protein A, *cusF* encoding Cu cation efflux system protein; *mrpA* and *nhaP* encoding Na^+^/H^+^ antiporter, *natB* encoding ABC transporter sodium permease; *yiip_fieF* encoding cation-efflux pump FieF, *zntA* encoding heavy metal translocating P-type ATPase. However, the relative abundances of several metal homeostasis genes were decreased in contaminated wells (Fig. 3D), including *silC* and *silP* encoding heavy metal RND efflux transporter; *chaA* encoding calcium/proton antiporter; *chrR* encoding chromate reductase; *cirA* encoding Colicin I receptor, *fhuE* encoding ferric-rhodotorulic acid outer membrane transporter, *dps* encoding DNA-binding ferritin-like protein; *kdpA* and *kup* encoding proteins in potassium transport system; *NiCoT* and *nikC* encoding nickel transport system proteins; *tehB, terC, terD, terZ*, and *terZD* encoding tellurite resistance protein (Fig. 3D).

### Stress response genes

Most osmotic stress genes, including *kdpE* encoding a transcriptional regulatory protein, *mtrA* encoding a DNA-binding response regulator, *ompR* encoding an osmolarity response regulator, *opuE* encoding osmoregulated proline transporter, and *proX* encoding glycine betaine transporter periplasmic subunit, were decreased in contaminated wells (Fig. 3E). Oxidative stress genes were decreased in contaminated wells, including *katA* encoding catalase that catalyzes the hydrogen peroxide and *soxR* encoding redox-sensitive transcriptional activator. Two oxygen limitation response genes, *narH* encoding the beta subunit of nitrate reductase and *narI* encoding the gamma subunit of nitrate reductase were increased in mid- and high-contaminated wells, which suggested microbial response to low dissolved oxygen concentration in mid- and high-contaminated wells (Fig. 3E).

### Organic pollutant degradation genes

The relative abundances of most organic pollutant degradation genes were decreased in mid- and high-contaminated wells (Fig. 3F), including *bphA* encoding biphenyl dioxygenase subunit alpha that catalyzes the oxygenation of biphenyl, *cmcI* encoding 3-carboxy-cis,cis-muconate cycloisomerase, *hbh* encoding 4-hydroxybenzoate hydroxylase that degrades aromatic compounds, *nagG* encoding salicylate 1-monooxygenase that catalyzes the decarboxylative hydroxylation of salicylate, *oxdB* encoding phenylacetaldoxime dehydratase that degrades styrene, *pobA* encoding 4-hydroxybenzoate 3-monooxygenase that catalyzes the hydroxylation of 4-hydroxybenzoate, *tfdA* encoding taurine catabolism dioxygenase that involves in taurine and hypotaurine metabolism.

### Electron transfer genes

The relative abundances of certain electron transfer genes were increased in high-contaminated wells (Fig. 3G), including C-type cytochrome genes encoding cytochrome c-type biogenesis protein CcmA and CcmF. The relative abundances of other electron transfer genes were decreased in low-contaminted wells, including some C-type cytochrome genes encoding cytochrome c class I.

### The linkages between microbial communities and environmental factors

To explore the relative importance of various factors in explaining microbial communities, we carried out partial least squares modeling (PLS) followed by variation partition analysis (VPA, Fig. 4A). Environmental variables and geographical distance explained considerable percentages of community variations for taxonomic compositions (R^2^ = 0.557, *p* = 0.001) and phylogenetic compositions (R^2^ = 0.679, *p* = 0.001), which were substantially lower than the explanatory power for functional compositions (R^2^ = 0.897, *p* = 0.001). In addition, environmental variables were more important than the geographical distance in explaining the composition variations.

**Fig. 4.**
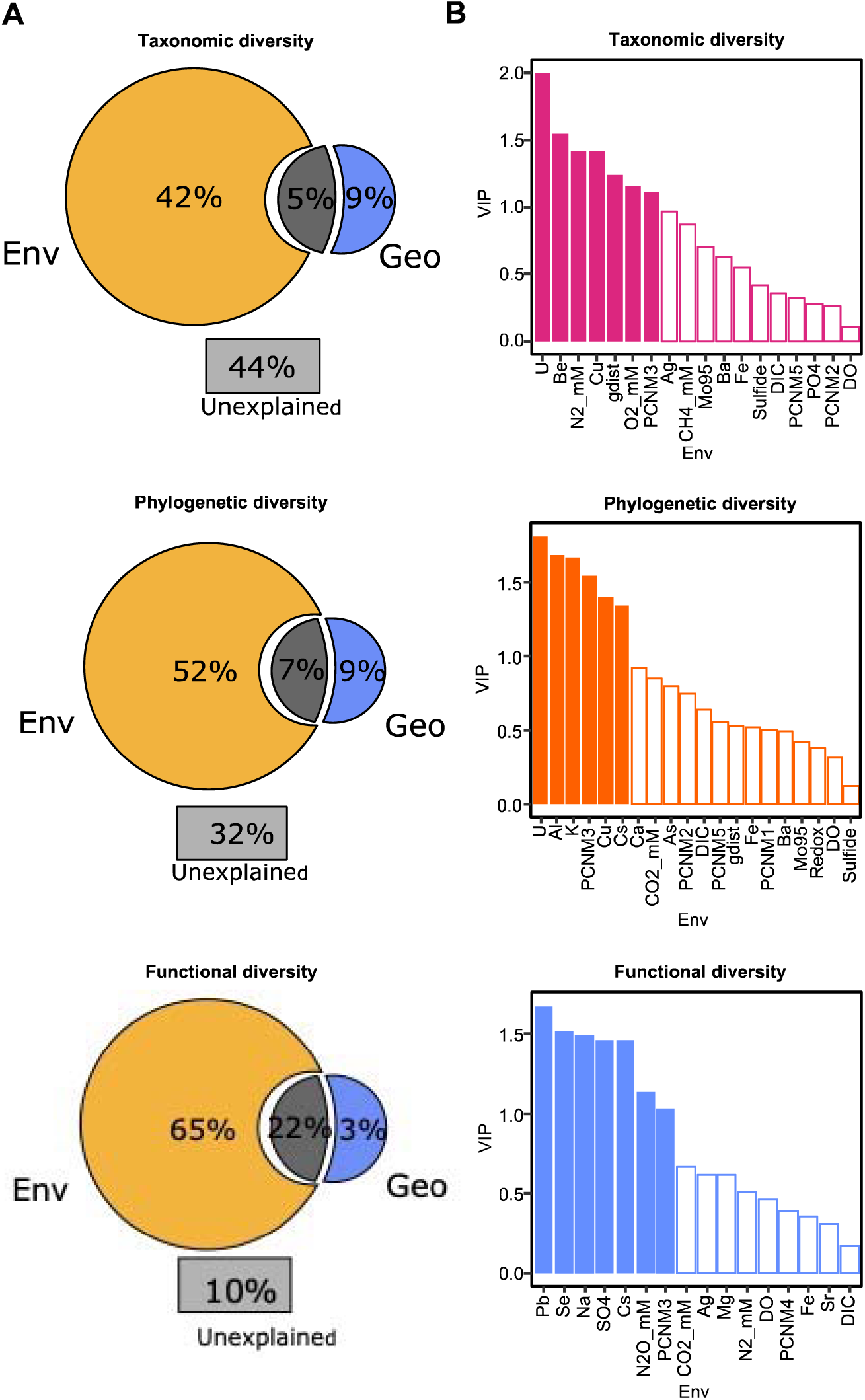
The linkage between aquifer microbial communities and environmental factors. (A) Variance partition analysis (VPA) showing relative contributions of geographical distance (Geo.) and environmental variables (Env.) to the different diversity indexes based on the PLS method. (B) Variable influence on projection (VIP) values based on the PLS model for different diversity indexes, where VIP value larger than 1 is filled, VIP value smaller than 1 is blank.

To evaluate the importance of individual environmental variables and geographic distance in PLS models, variable influence on projection (VIP) was calculated for taxonomic, phylogenetic, and functional diversity (Fig. 4B). DIC, DO, and Fe were important for all three dimensions of diversities (VIP > 0.10). Ba, Cu, U, and sulfide were important for taxonomic and phylogenetic diversities (VIP > 0.12), but not important for functional diversity. CO_2_ and Cs were important for phylogenetic and function diversities (VIP > 0.66), but not important for taxonomic diversity. Furthermore, geographic distance contributed more to taxonomic and phylogenetic diversities than functional diversity.

## DISCUSSION

Here, we explored three facets of microbial diversities, i.e., taxonomic, phylogenetic, and functional diversities along a broad spectrum of various contaminants. Consistent with the previous finding, (Rocca et al., 2018), we revealed a marked decrease in taxonomic and phylogenetic α-diversities in response to increasing contaminant levels (Fig. 1). In contrast, the reduction of functional α-diversities was much milder (Fig. 1). Our results also indicated an increase in functional composition heterogeneity correlating with environmental stressors, a pattern not mirrored in taxonomic and phylogenetic β-diversities (Fig. 2D). Consistently with findings in microbial communities associated with animal and plant hosts (Arnault et al., 2023; Zaneveld et al., 2017), aquifer microbial functional compositions in high-contaminated wells diverged more substantially than those in uncontaminated wells (Fig. 2), leading to increased dispersion in microbial community composition. Therefore, the Anna Karenina Principle is not limited to host-associated microbial communities, but is also applicable to free-living microbial communities and functional diversity. This suggests that the principle, indicative of microbial responses to environmental stress, might be a more widespread phenomenon.

The dispersion of environmental variables increases under contamination (Fig. S2). As a result, these variables become more dissimilar, leading to greater heterogeneity in functional diversity. Our study revealed a significant increase in the relative abundance of functional genes associated with denitrification and sulfite reduction in mid- and high-contaminated wells (Fig. 3B&3C), concurrent with the increased concentrations of nitrate and uranium. These findings align with previous research conducted at the same site (He et al., 2018; Xu et al., 2010; Zhang et al., 2015), which suggests that these functional genes are critical for heavy metal reduction.

Similar to a previous observation (Anantharaman et al., 2016), metabolic plasticity, involving various electron donors and acceptors, is a common trait in aquifer microorganisms. A wide metabolic repertoire is important in the face of the natural environmental perturbations that occur at the Oak Ridge site, where frequent storms and snows cause considerable water table fluctuations that move the oxic/anoxic interface. Consistent with earlier findings at our site (Carlson et al., 2019; Green et al., 2012), the denitrifying *Rhodanobacter*, known for its ability to immobilize U(VI) under aerobic conditions by forming intracellular uranium–phosphate complexes but not for U(VI) reduction (Green et al., 2012), was the dominant genus in the most contaminated wells characterized by low pH and high levels of nitrate and U concentrations (Table S1). Its dominance is likely due to its tolerance to NaCl and heavy metals (Prakash et al., 2021). A previous study has found that *Eremiobacterota* may be involved in Fe (II) oxidation (Grettenberger and Hamilton, 2021), which could explain the high relative abundance of the genus WPS-2 in high-contaminated wells with elevated ferrous levels (Table S2B).

*Sulfurifustis*, a genus of sulfur-oxidizing and glutathione-synthesizing bacteria, was enriched in high-contaminated wells (Table S2B) because glutathione synthesis serves as a mechanism for resisting cadmium toxicity (Liang et al., 2016). An ammonia-oxidizing archaeon named *Nitrosoarchaeum* was the most abundant genus in low-contaminated wells (Table S2B), which was also verified by more abundant *amoA* genes in those wells (Fig. 3B). In contrast, an ammonia-oxidizing bacterium named *GOUTA6* was the most abundant genus in mid-contaminated wells (Table S2B). Methanotrophs, including the genus *Candidatus Methylomirabilis,* can use methane to transform heavy metals (Karthikeyan et al., 2021), whose unique methane oxidation pathway requires both nitrate and methane (Versantvoort et al., 2018). Accordingly, we found that the abundance of *Candidatus Methylomirabilis* in mid-contaminated wells was characterized by the concentrations of nitrate and methane (Table S2B).

Functional redundancy may also explain the Anna Karenina Principle. Functional redundancy was termed as the coexistence of different species capable of performing the same biochemical functions, which could explain why the taxonomic and phylogenetic α-diversities decreased more noticeably than functional α-diversity under stress (Fig. 1). Functional diversity showed the lowest dispersion in uncontaminated wells because of the functional redundancy. In contrast, the dispersion values of functional diversity increased as the functional redundancy decreased with the contamination levels, whereas the dispersion values of taxonomic and phylogenetic diversities remained relatively unchanged (Fig. 2).

Both deterministic and stochastic processes contribute to the increased dissimilarity in stressed microbial communities, though the Anna Karenina Principle is mainly based on the important role of stochastic processes in disrupting normal community composition (Arnault et al., 2023; Zaneveld et al., 2017). The variability in key biogeochemical conditions (e.g., DIC, DO and Fe, Fig. 4B) emerged as important determinants of groundwater community compositions, which affected microbial fitness. A recent study revealed that stochastic processes, especially dispersal limitation, played an important role in shaping groundwater microbial communities (Ning et al., 2024), but the relative importance of stochastic processes decreased as contamination increased, which may explain why the Anna Karenina Principles was not observed in taxonomic and phylogenetic diversities. Other factors, including biotic interactions among community members and stochastic processes (e.g., ecological drift and dispersal limitation), could also play important roles in shaping community assembly (Chen et al., 2022); Zhou and Ning 2017).

## CONCLUSION

In this study, we assayed biological and geochemical diversities in a uranium-contaminated aquifer at Oak Ridge, TN, USA. Our results showed that environmental stressors have significant impacts on microbial diversity, particularly on taxonomic and phylogenetic diversities. The observed decrease in functional α-diversity was modest, indicating that the functional traits of the microbial communities had a better buffering capacity against environmental stress. Our results of functional heterogeneity (Fig. 2) explained the often low efficacy in treating *in-situ* groundwater contamination, which is costly and of large scale.

Understanding microbial functional responses in the stress environment is a central topic of microbial ecology. Therefore, our study is a useful asset for determining the critical factors linking community taxonomy to functions, which contribute to the development of accurate hydrogeochemical models that aid in assessing environmental treatments and evaluating risk management (King et al., 2017). The functional composition may be a sensitive and informative metric for evaluating the responses of microbial communities to environmental stress, which could inform the development of more effective and efficient bioremediation strategies for contaminated sites by providing a better understanding of the functional traits of microbial communities effective in degrading specific contaminants. Furthermore, our study demonstrated the importance of considering microbial functionality when evaluating the health of ecosystems, as the functional traits of microbial communities play a crucial role in maintaining ecosystem processes. Overall, our study contributes to a growing body of research that seeks to understand the functional response of microbial communities to environmental stress, and showed that microbial functionality should be taken in account in environmental management and risk assessment.

## MATERIALS AND METHODS

### Study site and sampling

We conducted this study at the Department of Energy’s (DOE) Oak Ridge FRC site in Oak Ridge, Tennessee. The groundwater at this location is tainted with various contaminants including radionuclides (such as uranium and technetium), nitrate, sulfide, and others, predominantly originating from the erstwhile S-3 waste disposal ponds. Groundwater samples were obtained from 12 representative wells along a gradient of various contaminants during winter 2012 and spring 2013 (Fig. S1): uncontaminated wells (UC) including FW300, FW301, and FW303 (FW305 for metagenome shotgun sequencing due to FW303 was not easy to access); low-contaminated wells (LC) including GW199, GW715, and GW928 with nitrate concentrations less than 2 mg/L, uranium concentrations less than 0.01 mg/L, and neutral pH (6.5-7.2); mid-contaminated wells (MC) including FW215, FW602, and DP16D with nitrate concentrations between 5.5 to 1471 mg/L, uranium concentrations between 0.1 to 1.5 mg/L, and neutral pH (6.5-6.8); high-contaminated wells (HC) including FW104, FW106, FW021 with nitrate concentrations between 2692 to 11648 mg/L, uranium concentrations between 3.8 to 55 mg/L, and low pH (3-5.2).

### Geophysical and geochemical analyses

The groundwater properties of temperature, pH, dissolved oxygen (DO), conductivity, and redox were measured by an In-Situ Troll 9500 system (In-Situ Inc., CO, USA). U.S. EPA methylene blue method (Hach; EPA Method 8131) and the 1,10-phenanthroline method (Hach; EPA Method 8146) were used to measure sulfide and ferrous iron concentrations, respectively. The concentrations of dissolved gases (N_2_, O_2_, CO_2_, CH_4_, and N_2_O) were determined using an SRI 8610C gas chromatograph (GC) with argon as the carrier gas. The method was derived from EPA RSK-175 and United States Geological Survey (USGS) Reston Chlorofluorocarbon Laboratory protocols. Concentrations of dissolved organic carbon (DOC) and dissolved inorganic carbon (DIC) were ascertained using a Shimadzu TOC-V CSH analyzer (Tokyo, Japan), following the EPA Method 415.1. Anion concentrations, including bromide, chloride, nitrate, phosphate, and sulfate, were quantified using a Dionex 2100 system equipped with an AS9 column and a carbonate eluent, in accordance with U.S. EPA Methods 300.1 and 317.0. The levels of metals and trace elements present in the groundwater were assessed using an inductively coupled plasma/mass spectrometry (ICP-MS) instrument (Elan 6100), employing a technique akin to EPA Method 200.7.

### Amplicon and metagenomic sequencing

The DNA extraction method was described in a previous study (Smith et al., 2015). The phasing amplicon sequencing (PAS) method (Wu et al., 2015) was used to amplify the V4 region of 16S rRNA genes and the samples were sequenced on an Illumina MiSeq platform. The primers are 515F (5’-GTGCCAGCMGCCGCGGTAA-3’) and 806R (5’-GGACTACHVGGGTWTCTAAT-3’). The sequencing data were processed by Qiime2 (version 2019.7). After barcode and primer sequences were trimmed with zero maximum error, sequencing data were processed by DADA2 to identify exact amplicon sequence variants (ASV). The ASVs were identified taxonomically based on the silva-132-99-515-806-nb-classifier. The ASV sequences were then used to build a phylogenetic tree by FastTree (Price et al., 2009; Price et al., 2010).

We used the KAPA Hyper Prep Kit (KR0961) to construct the metagenomic sequencing libraries following the manufacturer’s instructions, and the samples were sequenced using an Illumina HiSeq 3000 sequencer. The read-based metagenomic data analysis was performed using an internal pipeline (http://iegst1.rccc.ou.edu:8080/ecofunmap/) following the guidelines from Shi et al., 2022. For assembly-based analysis, metagenomic reads were preprocessed using BBTools for removing adaptor, trimming and filtering reads, and sequencing error correction (Bushnell 2014). The pre-processed reads were assembled with Metaspades (Nurk et al., 2017). Genes were predicted from scaffolds > 1kbp using the Prodigal (Hyatt et al., 2010). The gene abundance was estimated as TPM (Zhao et al., 2020). Genes functions were annotated using the Kofamsan (Kanehisa et al., 2020). Species-level quantitative taxonomic profiling was performed using MetaPhlAn4 (Blanco-Míguez et al., 2023).

### Statistical analyses

We used richness to represent microbial α-diversity and analysis of variance (ANOVA) to compare the difference between each group. Non-metric multidimensional (NMDS) and permutation test of multivariate homogeneity of groups dispersions (Anderson et al. 2006) were used for microbial β-diversity, taxonomic and functional β-diversity were measured by Bray-Curtis dissimilarity (Bray and Curtis 1957) and phylogenetic β-diversity were measured by UniFrac (Lozupone et al 2007). We used permutation test of multivariate homogeneity of groups dispersions (Anderson et al. 2006) based on the Euclidean distance for environmental variables. The statistical significance of the effects of contaminations on β-diversity was tested by multi response permutation procedure (MRPP) (Warton et al 2012), permutational multivariate analysis of variance (Adonis) (Anderson 2017), and analysis of similarities (Anosim) (Clarke 1993, Warton et al 2012) by using function ‘mrpp’, ‘adonis’, and ‘anosim’ in R package ‘vegan’, respectively (Oksanen et al. 2020). Microbial γ-diversity was shown for richness. The response ratio was calculated using an internal R packcage ‘ieggr’ based on 95% confidence intervals. We used a partial least squares (PLS) model to detect the relationships between environmental variables and geographical distance for each diversity index. Basically, each optimal PLS model is selected through a forward selection process from all factors that could influence the dependent variable. This selection is based on predictive performance, taking into account the proportion of variation explained (R²Y) and the statistical significance of the model (P values for R²Y and Q²Y less than 0.05). Notably, a significant Q²Y aids in preventing overfitting of the model. To visualize the relevant associations, we used the variance partition analysis (VPA) and the software Inkscape 1.3 (https://inkscape.org/). The PLS-related analysis was performed using the R package “ropls” (Thevenot et al. 2015).

## Declarations

### Ethics approval and consent to participate

Not applicable.

### Consent for publication

Not applicable.

### Funding

This work is supported by the US Department of Energy, Office of Science, Biological and Environmental Research Program under contract number DE-AC02-05CH11231 to Lawrence Berkeley National Lanoratory.

### Availability of data and materials

The 16S rRNA gene amplicon sequences were submitted to NCBI database SRA under the project PRJNA514085 with accession SRR8427255. The shotgun metagenomic sequences were submitted to NCBI database SRA under the project PRJNA513876 with accession SRR8426587 - SRR8426598.

### Competing interests

The authors declare that they have no competing interests.

### Authors’ contributions

All authors contributed intellectual input and assistance to this study and the paper preparation. J. Z.. conceived the research question. Z. H., M. W. W. A., M. W. F., E. J. A., T. C. H., P. D. A., A. P. A., and J. Z. designed and organized the experiment. P. Z., A. M. R., J. D. V. N., and D. C. J. collected or generated the data. Y.F., D. W., J. X. Y. and J. P. M. intergrated the data and performed statistical analyses with the assistance of D.N. and C. P.; Y.F. and J. X. Y. wrote the paper with inputs from D.N. and J.Z.

## Acknowledgements

This study by ENIGMA (Ecosystems and Networks Integrated with Genes and Molecular Assemblies; http://enigma.lbl.gov), a Science Focus Area Program at Lawrence Berkeley National Laboratory, is based on work supported by the US Department of Energy, Office of Science, Biological and Environmental Research Program under contract number DE-AC02-05CH11231 to Lawrence Berkeley National Lanoratory.

---

**Table S1.** Aquifer environmental variables in in uncontaminated wells (UC), low-contaminated wells (LC), mid-contaminated wells (MC), and high-contaminated wells (HC).

|  | UC | LC | MC | HC |
| --- | --- | --- | --- | --- |
| Conductivity<br>($\mu\text{S}/\text{cm}$) | 242.53±175.75<sup>b</sup> | 452±183.92<sup>b</sup> | 1000.8±609.1<sup>b</sup> | 9830.33±3316.99<sup>a</sup> |
| pH | 6.43±0.94<sup>b</sup> | 6.62±0.08<sup>b</sup> | 6.58±0.1<sup>b</sup> | 4.06±1<sup>a</sup> |
| Redox (mV) | 42.33±151.79<sup>ab</sup> | 474.33±287.25<sup>ab</sup> | 32.67±78.01<sup>a</sup> | 332.67±129.36<sup>b</sup> |
| DO (mg/L) | 1.87±1.65<sup>a</sup> | 2.06±2.96<sup>a</sup> | 0.27±0.14<sup>a</sup> | 0.54±0.55<sup>a</sup> |
| DIC (mg/L) | 34.21±17.03<sup>a</sup> | 54.29±28.13<sup>a</sup> | 74.96±20.17<sup>a</sup> | 60.81±49.77<sup>a</sup> |
| DOC (mg/L) | 15.24±25.38<sup>ab</sup> | 0.86±0.44<sup>a</sup> | 1.95±0.37<sup>b</sup> | 20.56±23.65<sup>ab</sup> |
| O<sub>2</sub> (mM) | 0.25±0.12<sup>a</sup> | 0.5±0.25<sup>a</sup> | 0.23±0.09<sup>a</sup> | 0.17±0.06<sup>a</sup> |
| CH<sub>4</sub> (mM) | 0.23±0.23<sup>a</sup> | 0.02±0.02<sup>a</sup> | 0.02±0.006<sup>a</sup> | 0 |
| CO<sub>2</sub> (mM) | 12.05±18.25<sup>ab</sup> | 3.49±1.41<sup>b</sup> | 4.4±1.57<sup>b</sup> | 15.06±5.84<sup>a</sup> |
| N<sub>2</sub> (mM) | 1.42±0.45<sup>ab</sup> | 2.48±0.95<sup>a</sup> | 1.11±0.39<sup>ab</sup> | 0.69±0.24<sup>b</sup> |
| N<sub>2</sub>O (mM) | 0.11±0.11<sup>a</sup> | 0 | 0.1±0.1<sup>a</sup> | 0.88±0.27<sup>b</sup> |
| S<sup>2-</sup> (mg/L) | 0.026±0.026<sup>a</sup> | 0.002±0.001<sup>a</sup> | 0.001±0.001<sup>a</sup> | 0.002±0.002<sup>a</sup> |
| Cl<sup>-</sup> (mg/L) | 1.63±0.49<sup>c</sup> | 12.35±15.65<sup>bc</sup> | 21.04±4.72<sup>b</sup> | 238.36±83.1<sup>a</sup> |
| NO<sub>3</sub><sup>-</sup> (mg/L) | 1.36±2<sup>b</sup> | 1.16±1.87<sup>b</sup> | 321.03±434.46<sup>ab</sup> | 5422.55±3285.46<sup>a</sup> |
| PO<sub>4</sub><sup>-</sup> (mg/L) | ND | 0.08±0.13 | ND | ND |
| SO<sub>4</sub><sup>2-</sup> (mg/L) | 7.16±0.91<sup>a</sup> | 15±5.16<sup>a</sup> | 49.48±36.62<sup>a</sup> | 799.51±1101.18<sup>a</sup> |
| NO<sub>3</sub>_sup (mM) | ND | 0.02±0.04<sup>a</sup> | 3.25±5.56<sup>a</sup> | 38.65±33.63<sup>a</sup> |
| NO<sub>2</sub>_sup ($\mu\text{M}$) | ND | ND | 19.51±24.26<sup>a</sup> | 13.61±4.49<sup>a</sup> |
| Ferrous (mg/L) | 0.18±0.28<sup>a</sup> | 0.33±0.45<sup>a</sup> | 0.56±0.79<sup>a</sup> | 1.08±1.07<sup>a</sup> |
| K (mg/L) | 1.3±0.29<sup>a</sup> | 3.09±1.59<sup>ab</sup> | 4.8±1.38<sup>b</sup> | 111.38±95.81<sup>ab</sup> |
| Na (mg/L) | ND | ND | 16.35±14.93<sup>a</sup> | 508.44±315<sup>a</sup> |
| Ca (mg/L) | 41.29±33.12<sup>a</sup> | 69.19±26.08<sup>a</sup> | 704.8±995.72<sup>a</sup> | 5463.17±6075.62<sup>a</sup> |
| Mg (mg/L) | ND | ND | 9.96±17.25<sup>a</sup> | 117.05±81.56<sup>a</sup> |
| Al (mg/L) | 0.01±0.02<sup>a</sup> | 0.6±1.01<sup>a</sup> | 1.15±1.99<sup>a</sup> | 76.27±61.1<sup>a</sup> |
| Ba (mg/L) | 0.07±0.02<sup>a</sup> | 0.1±0.01<sup>a</sup> | 0.33±0.26<sup>a</sup> | 0.82±1.09<sup>a</sup> |
| Ag (mg/L) | 0.005±0.004<sup>b</sup> | 0.01±0.0007<sup>ab</sup> | 0.007±0.006<sup>ab</sup> | 0.018±0.006<sup>a</sup> |
| As (mg/L) | ND | 0.0006±0.001<sup>a</sup> | ND | 0.003±0.005<sup>a</sup> |
| Be (mg/L) | ND | 0.006±0.01<sup>a</sup> | ND | 0.06±0.02<sup>b</sup> |
| Bi (mg/L) | ND | 0.001±0.005 | ND | ND |
| Cd (mg/L) | ND | 0.002±0.001<sup>a</sup> | 0.003±0.006<sup>a</sup> | 0.142±0.03<sup>b</sup> |
| Co (mg/L) | ND | 0.01±0.02<sup>a</sup> | 0.02±0.03<sup>a</sup> | 0.62±0.55<sup>a</sup> |
| Cr (mg/L) | ND | 0.004±0.002<sup>a</sup> | 0.002±0.003<sup>a</sup> | 0.13±0.22<sup>a</sup> |
| Cs (mg/L) | ND | 0.003±0.005<sup>a</sup> | ND | 0.011±0.01<sup>a</sup> |
| Cu (mg/L) | ND | 0.014±0.002<sup>a</sup> | ND | 0.32±0.42<sup>a</sup> |
| Zn (mg/L) | 0.06±0.01<sup>a</sup> | 0.06±0.02<sup>a</sup> | 0.05±0.02<sup>a</sup> | 0.73±0.47<sup>a</sup> |
| Fe (mg/L) | 0.02±0.04<sup>a</sup> | 0.76±0.41<sup>a</sup> | 0.64±1.12<sup>a</sup> | 1.08±1.84<sup>a</sup> |
| Ga (mg/L) | ND | 0.005±0.004<sup>a</sup> | 0.011±0.01<sup>a</sup> | 0.03±0.02<sup>a</sup> |
| Mn (mg/L) | 0.07±0.11<sup>a</sup> | 1.58±2.48<sup>a</sup> | 3.38±5.21<sup>a</sup> | 95±54.18<sup>b</sup> |
| Li (mg/L) | ND | 0.02±0.01<sup>a</sup> | 0.016±0.03<sup>a</sup> | 0.75±1.04<sup>a</sup> |
| Ni (mg/L) | ND | 0.051±0.09<sup>a</sup> | 0.081±0.14<sup>a</sup> | 4.52±3.15<sup>a</sup> |
| Pb (mg/L) | ND | 0.0008±0.0007<sup>a</sup> | ND | 0.013±0.016<sup>a</sup> |
| Se (mg/L) | ND | ND | ND | 0.012±0.012 |
| Sr (mg/L) | 0.15±0.04<sup>a</sup> | 0.10±0.11<sup>a</sup> | 0.52±0.23<sup>a</sup> | 1.7±1.52<sup>a</sup> |
| U (mg/L) | 0.07±0.12<sup>a</sup> | 0.004±0.004<sup>b</sup> | 0.75±0.7<sup>ab</sup> | 9.79±6.47<sup>ab</sup> |
| Mo<sup>95</sup>_sup (μM) | 0.002±0.003<sup>a</sup> | 0.001±0.0009<sup>a</sup> | 0.01±0.02<sup>a</sup> | 0.004±0.007<sup>a</sup> |

<sup>a</sup>Letters represent the difference between groups. The difference between groups was determined by ANOVA. Significant difference level: $p < 0.05$ . ND: Not detected; sup: concentration in the supernatant of each sample.

**Fig. S1.**
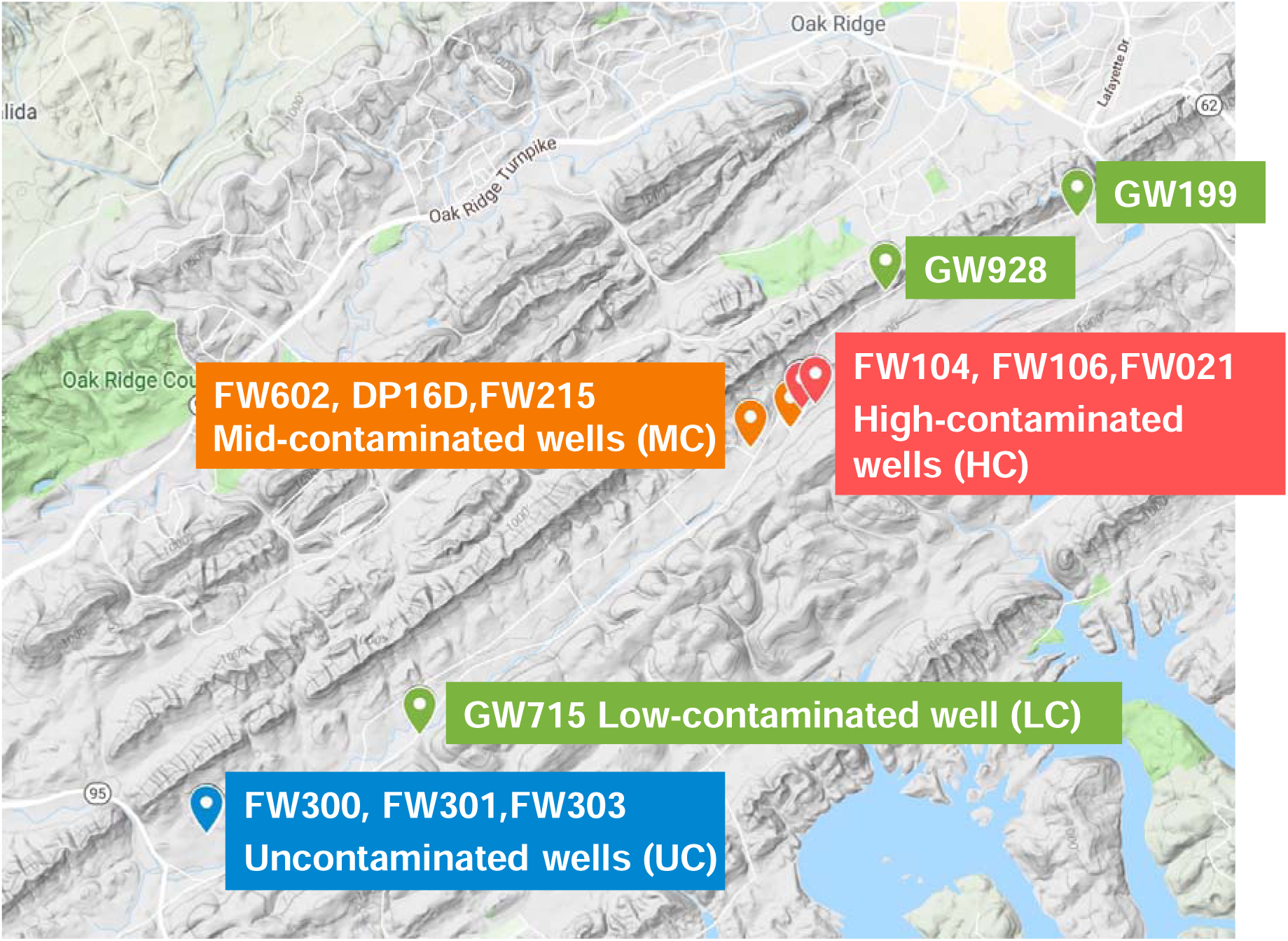
Geographical location of the study sites. Aquifer samples consist of uncontaminated wells (UC) FW300, FW301, FW303, and FW305; low-contaminated wells (LC) GW199, GW715, and GW928; mid-contaminated wells (MC) FW215, FW602, and DP16D; high-contaminated wells (HC) FW104, FW106, and FW021.

**Fig. S2.**
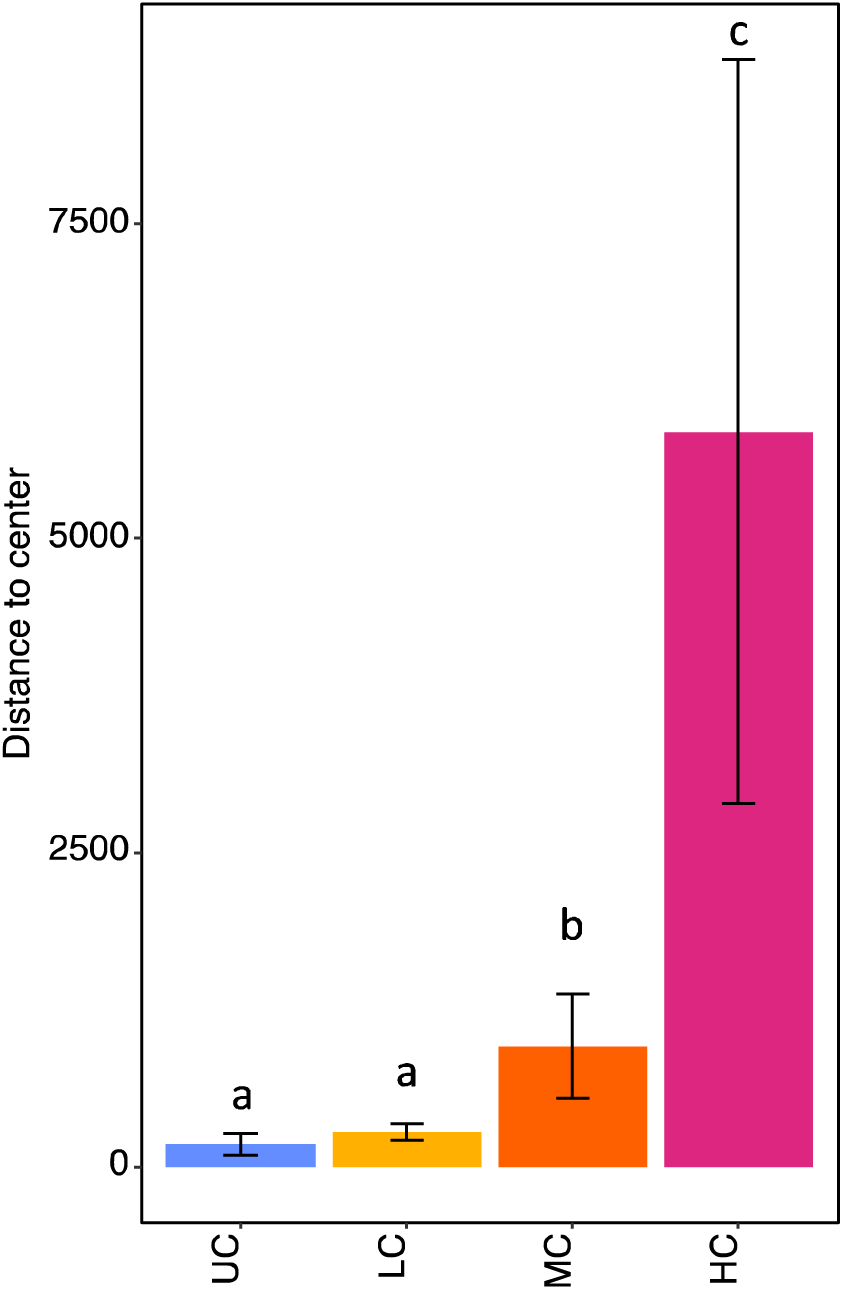
Dispersion analysis of environmental variables based on Euclidean distance. Letters represent the difference between groups which was determined by ANOVA (significant difference level: p < 0.05).

**Fig. S3.**
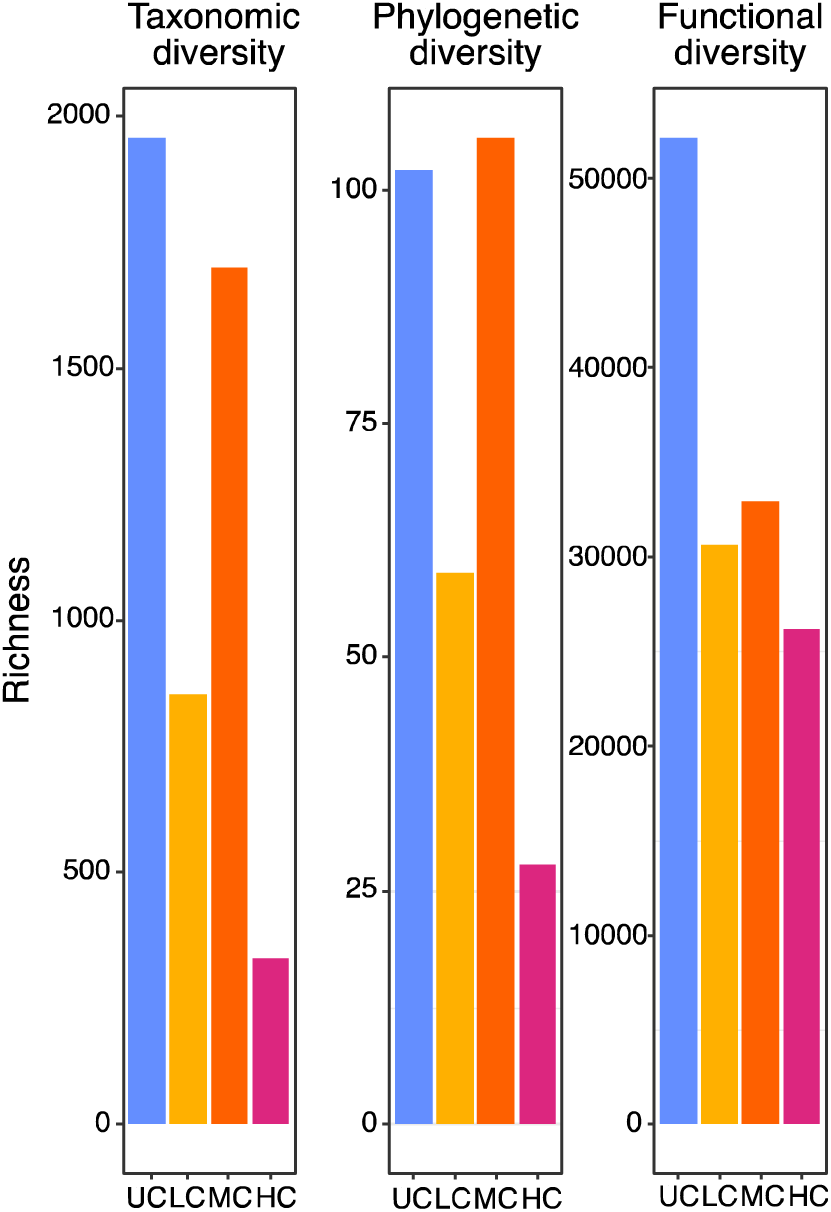
Microbial γ-diversity of 12 groundwater samples calculated by richness.

**Table S2.** Taxonomic composition of microbial communities at phylum (A) and genus (B) level under different treatments.

| A.Category at Phylum level | UC | LC | MC | HC |
| --- | --- | --- | --- | --- |
| | Avergae relative abundance ± standard deviation (%) | | | |
| Gammaproteobacteria | 42.86±19.18<sup>a</sup> | 27.21±15.73<sup>a</sup> | 23.45±1.90<sup>a</sup> | 73.06±22.44<sup>a</sup> |
| Alphaproteobacteria | 12.97±7.44<sup>b</sup> | 18.39±11.88<sup>a</sup> | 3.36±0.41<sup>cd</sup> | 1.25±1.89<sup>b</sup> |
| Verrucomicrobiota | 4.76±3.14<sup>b</sup> | 3.97±2.44<sup>a</sup> | 16.29±4.36<sup>ab</sup> | 4.54±7.74<sup>b</sup> |
| Bacteroidota | 3.22±3.10<sup>b</sup> | 11.72±6.57<sup>a</sup> | 5.15±2.62<sup>bc</sup> | 1.64±1.37<sup>b</sup> |
| Acidobacteriota | 2.20±1.08<sup>b</sup> | 0.83±0.56<sup>b</sup> | 6.33±5.11<sup>bc</sup> | 5.47±7.40<sup>b</sup> |
| WPS-2 | 0.22±0.23<sup>b</sup> | 0.07±0.08<sup>b</sup> | 0.54±0.74<sup>c</sup> | 12.45±21.56<sup>b</sup> |
| Nitrospirota | 2.54±2.75<sup>b</sup> | 6.72±6.98<sup>a</sup> | 2.57±1.90<sup>c</sup> | 0.06±0.05<sup>b</sup> |
| Patescibacteria | 3.43±2.01<sup>b</sup> | 0.41±0.30<sup>b</sup> | 7.02±4.53<sup>bc</sup> | 0.31±0.51<sup>b</sup> |
| Bdellovibrionota | 8.39±10.41<sup>b</sup> | 0.84±1.22<sup>b</sup> | 0.96±1.01<sup>c</sup> | 0.00±0.00<sup>b</sup> |
| Myxococcota | 1.13±0.97<sup>b</sup> | 1.92±2.55<sup>b</sup> | 5.97±2.90<sup>bc</sup> | 0.02±0.02<sup>b</sup> |
| Methylomirabilota | 0.70±0.89<sup>b</sup> | 0.22±0.30<sup>b</sup> | 7.04±7.13<sup>bc</sup> | 0.01±0.01<sup>b</sup> |
| Elusimicrobiota | 4.06±2.44<sup>b</sup> | 0.81±1.10<sup>b</sup> | 1.58±1.37<sup>c</sup> | 0.01±0.01<sup>b</sup> |
| Campilobacterota | 6.06±10.32<sup>b</sup> | 0.00±0.00<sup>b</sup> | 0.02±0.01<sup>c</sup> | 0.00±0.00<sup>b</sup> |
| Chloroflexi | 0.46±0.41<sup>b</sup> | 0.70±0.98<sup>b</sup> | 4.01±5.90<sup>cd</sup> | 0.11±0.15<sup>b</sup> |
| DTB120 | 0.12±0.18<sup>b</sup> | 0.04±0.06<sup>b</sup> | 3.73±5.58<sup>cd</sup> | 0.12±0.21<sup>b</sup> |
| Actinobacteriota | 0.60±0.51<sup>b</sup> | 2.04±1.81<sup>a</sup> | 0.72±0.42<sup>c</sup> | 0.12±0.08<sup>b</sup> |
| Spirochaetota | 0.09±0.06<sup>b</sup> | 0.01±0.02<sup>b</sup> | 2.66±2.62<sup>c</sup> | 0.14±0.24<sup>b</sup> |
| Crenarchaeota | 0.20±0.05<sup>b</sup> | 22.93±30.70<sup>a</sup> | 0.17±0.18<sup>c</sup> | 0.00±0.00<sup>b</sup> |
| Others | 6.00±2.45<sup>b</sup> | 1.16±0.57<sup>b</sup> | 8.45±3.46<sup>bc</sup> | 0.69±0.78<sup>b</sup> |

| B.Category at Genus level | UC | LC | MC | HC |
| --- | --- | --- | --- | --- |
| | Avergae relative abundance ± standard deviation (%) | | | |
| Rhodanobacter | 0.01±0.02<sup>c</sup> | 0.02±0.03<sup>b</sup> | 0.86±0.89<sup>c</sup> | 37.50±35.73<sup>a</sup> |
| Nitrosarchaeum | 0.06±0.03<sup>c</sup> | 19.18±33.14<sup>ab</sup> | 0.14±0.20<sup>c</sup> | 0.00±0.00<sup>b</sup> |
| Candidatus_Omnitrophus | 1.14±1.81<sup>c</sup> | 3.14±2.60<sup>b</sup> | 10.05±8.20<sup>bc</sup> | 3.93±6.72<sup>ab</sup> |
| WPS-2 | 0.22±0.23<sup>c</sup> | 0.07±0.08<sup>b</sup> | 0.54±0.74<sup>c</sup> | 12.45±21.56<sup>ab</sup> |
| Sulfuritalea | 1.72±0.99<sup>c</sup> | 9.85±8.40<sup>ab</sup> | 0.07±0.12<sup>c</sup> | 0.00±0.00<sup>b</sup> |
| Sulfurifustis | 0.44±0.56<sup>c</sup> | 0.10±0.09<sup>b</sup> | 1.56±1.57<sup>bc</sup> | 9.51±16.46<sup>ab</sup> |
| Chujaibacter | 0.00±0.00<sup>c</sup> | 0.00±0.00<sup>b</sup> | 1.01±1.09<sup>c</sup> | 7.52±13.03<sup>ab</sup> |
| Reyranella | 0.84±1.08<sup>c</sup> | 6.25±7.19<sup>b</sup> | 0.13±0.13<sup>c</sup> | 0.48±0.82<sup>b</sup> |
| GOUTA6 | 0.90±1.19<sup>c</sup> | 0.18±0.32<sup>b</sup> | 5.47±1.33<sup>bc</sup> | 0.40±0.70<sup>b</sup> |
| Candidatus_Methylomirabilis | 0.00±0.00<sup>c</sup> | 0.03±0.04<sup>b</sup> | 6.79±7.08<sup>bc</sup> | 0.01±0.01<sup>b</sup> |
| Nitrospira | 1.24±1.19<sup>c</sup> | 5.35±5.37<sup>b</sup> | 0.23±0.31<sup>c</sup> | 0.00±0.00<sup>b</sup> |
| 0319-6G20 | 5.93±7.67<sup>bc</sup> | 0.22±0.35<sup>b</sup> | 0.58±0.49<sup>c</sup> | 0.00±0.00<sup>b</sup> |
| bacteriap25 | 0.55±0.60<sup>c</sup> | 0.55±0.86<sup>b</sup> | 5.50±2.94<sup>bc</sup> | 0.00±0.00<sup>b</sup> |
| Brevundimonas | 1.51±1.76<sup>c</sup> | 4.79±5.84<sup>b</sup> | 0.04±0.07<sup>c</sup> | 0.00±0.00<sup>b</sup> |
| Sulfuricurvum | 5.90±10.04<sup>bc</sup> | 0.01±0.01<sup>b</sup> | 0.00±0.01<sup>c</sup> | 0.00±0.00<sup>b</sup> |
| Omnitrophales | 0.96±0.60<sup>c</sup> | 0.28±0.41<sup>b</sup> | 4.34±5.83<sup>bc</sup> | 0.00±0.00<sup>b</sup> |
| Sediminibacterium | 1.49±2.21<sup>c</sup> | 3.45±2.04<sup>b</sup> | 0.00±0.01<sup>c</sup> | 0.22±0.38<sup>b</sup> |
| Others | 46.78±4.31<sup>a</sup> | 34.94±8.97<sup>a</sup> | 42.62±7.12<sup>a</sup> | 14.90±14.43<sup>ab</sup> |
| unclassified | 18.24±12.41<sup>b</sup> | 7.18±4.16<sup>b</sup> | 8.34±2.45<sup>bc</sup> | 11.24±5.96<sup>ab</sup> |
| uncultured | 12.08±8.58<sup>bc</sup> | 4.40±2.60<sup>b</sup> | 11.71±3.03<sup>b</sup> | 1.83±1.50<sup>b</sup> |

<sup>a</sup>Letters represent the difference between groups. The difference between groups was determined by ANOVA. Significant difference level: $p < 0.05$ .

**Fig. S4.**
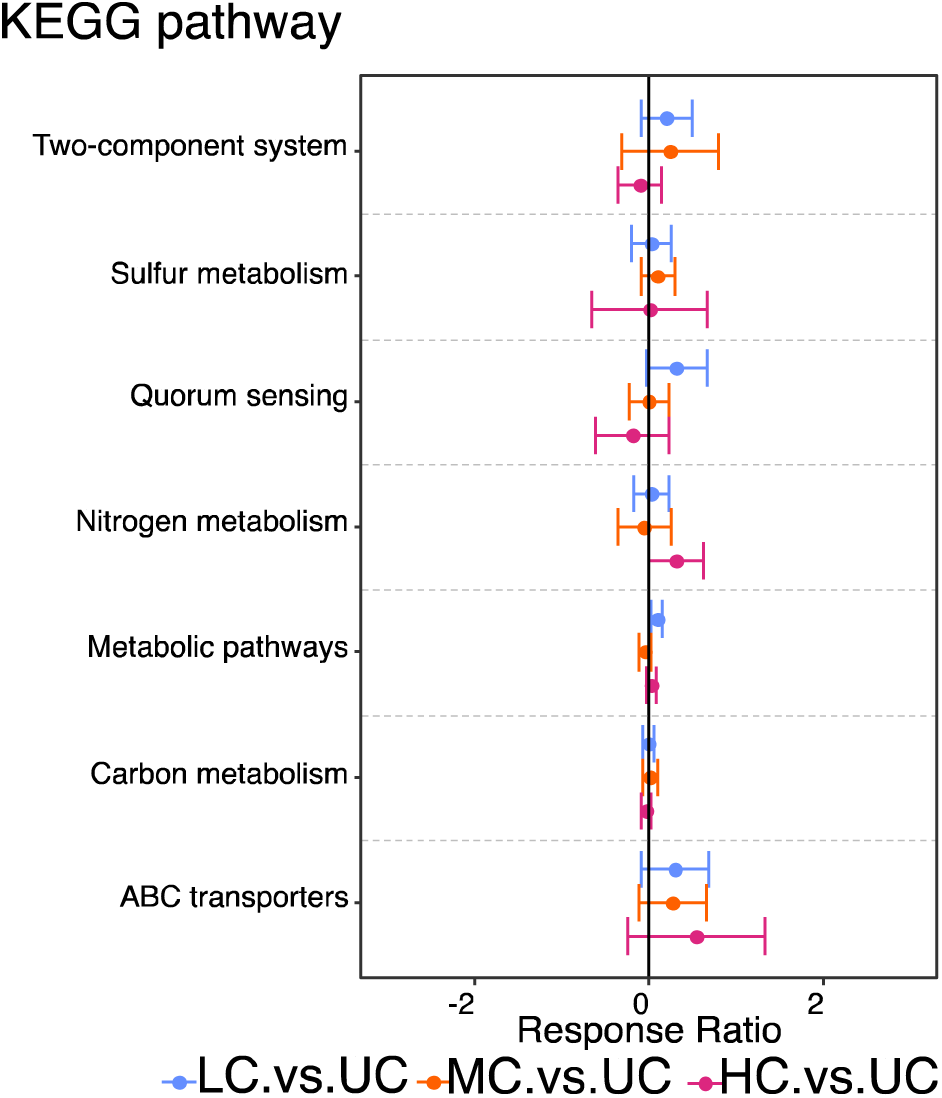
Differences in assembly-based shotgun metagenomic data between contaminated wells and uncontaminated wells, reflected by response ratios of annotated KEGG pathway.

## Notes

### Competing Interest Statement

The authors have declared no competing interest.

## REFERENCES

Anantharaman, K., Brown, C.T., Hug, L.A., Sharon, I., Castelle, C.J., Probst, A.J., Thomas, B.C., Singh, A., Wilkins, M.J., Karaoz, U., Brodie, E.L., Williams, K.H., Hubbard, S.S. and Banfield, J.F. 2016. Thousands of microbial genomes shed light on interconnected biogeochemical processes in an aquifer system. Nat Commun 7, 13219.

Anderson, M.J. 2006. Distance-based tests for homogeneity of multivariate dispersions. Biometrics 62(1), 245–253.

Anderson, M.J. 2017. Wiley StatsRef: Statistics Reference Online, pp. 1–15

Arnault, G., Mony, C. and Vandenkoornhuyse, P. 2023. Plant microbiota dysbiosis and the Anna Karenina Principle. Trends Plant Sci 28(1), 18–30.

Blanco-Miguez, A., Beghini, F., Cumbo, F., McIver, L.J., Thompson, K.N., Zolfo, M., Manghi, P., Dubois, L., Huang, K.D., Thomas, A.M., Nickols, W.A., Piccinno, G., Piperni, E., Puncochar, M., Valles-Colomer, M., Tett, A., Giordano, F., Davies, R., Wolf, J., Berry, S.E., Spector, T.D., Franzosa, E.A., Pasolli, E., Asnicar, F., Huttenhower, C. and Segata, N. 2023. Extending and improving metagenomic taxonomic profiling with uncharacterized species using MetaPhlAn 4. Nat Biotechnol 41(11), 1633–1644.

Bushnell, B., BBTools software packag. e, 2014.

Bray, J. R. and Curtis, J. T. 1957. An Ordination of the Upland Forest Communities of Southern Wisconsin. Ecological Monographs 27(4), 325–349.

Carlson, H.K., Price, M.N., Callaghan, M., Aaring, A., Chakraborty, R., Liu, H., Kuehl, J.V., Arkin, A.P. and Deutschbauer, A.M. 2019. The selective pressures on the microbial community in a metal-contaminated aquifer. ISME J 13(4), 937–949.

Chen, H., Chen, Z., Chu, X., Deng, Y., Qing, S., Sun, C., Wang, Q., Zhou, H., Cheng, H., Zhan, W. and Wang, Y. 2022. Temperature mediated the balance between stochastic and deterministic processes and reoccurrence of microbial community during treating aniline wastewater. Water Res 221, 118741.

Chen, J., He, F., Zhang, X., Sun, X., Zheng, J. and Zheng, J. 2014. Heavy metal pollution decreases microbial abundance, diversity and activity within particle-size fractions of a paddy soil. FEMS Microbiol Ecol 87(1), 164–181.

Clarke, K.R. 1993. Non-parametric multivariate analyses of changes in community structure. Australian Journal of Ecology 18(1), 117–143.

Deng, Y., Fu, S., Sarkodie, E.K., Zhang, S., Jiang, L., Liang, Y., Yin, H., Bai, L., Liu, X., Liu, H. and Jiang, H. 2022. Ecological responses of bacterial assembly and functions to steep Cd gradient in a typical Cd-contaminated farmland ecosystem. Ecotoxicol Environ Saf 229, 113067.

Ernakovich, J.G., Lynch, L.M., Brewer, P.E., Calderon, F.J. and Wallenstein, M.D. 2017. Redox and temperature-sensitive changes in microbial communities and soil chemistry dictate greenhouse gas loss from thawed permafrost. Biogeochemistry 134(1-2), 183–200.

Fierer, N., Ladau, J., Clemente, J.C., Leff, J.W., Owens, S.M., Pollard, K.S., Knight, R., Gilbert, J.A. and McCulley, R.L. 2013. Reconstructing the microbial diversity and function of pre-agricultural tallgrass prairie soils in the United States. Science 342(6158), 621–624.

Galand, P.E., Pereira, O., Hochart, C., Auguet, J.C. and Debroas, D. 2018. A strong link between marine microbial community composition and function challenges the idea of functional redundancy. ISME J 12(10), 2470–2478.

Gao, Q., Dong, Q., Wu, L., Yang, Y., Hale, L., Qin, Z., Xie, C., Zhang, Q., Van Nostrand, J.D. and Zhou, J. 2020. Environmental antibiotics drives the genetic functions of resistome dynamics. Environ Int 135, 105398.

Green, S.J., Prakash, O., Jasrotia, P., Overholt, W.A., Cardenas, E., Hubbard, D., Tiedje, J.M., Watson, D.B., Schadt, C.W., Brooks, S.C. and Kostka, J.E. 2012. Denitrifying bacteria from the genus Rhodanobacter dominate bacterial communities in the highly contaminated subsurface of a nuclear legacy waste site. Appl Environ Microbiol 78(4), 1039–1047.

Grettenberger, C.L. and Hamilton, T.L. 2021. Metagenome-Assembled Genomes of Novel Taxa from an Acid Mine Drainage Environment. Appl Environ Microbiol 87(17), e0077221.

Handley, K.M., Bartels, D., O’Loughlin, E.J., Williams, K.H., Trimble, W.L., Skinner, K., Gilbert, J.A., Desai, N., Glass, E.M., Paczian, T., Wilke, A., Antonopoulos, D., Kemner, K.M. and Meyer, F. 2014. The complete genome sequence for putative H(2)- and S-oxidizer Candidatus Sulfuricurvum sp., assembled de novo from an aquifer-derived metagenome. Environ Microbiol 16(11), 3443–3462.

He, Z., Zhang, P., Wu, L., Rocha, A.M., Tu, Q., Shi, Z., Wu, B., Qin, Y., Wang, J., Yan, Q., Curtis, D., Ning, D., Van Nostrand, J.D., Wu, L., Yang, Y., Elias, D.A., Watson, D.B., Adams, M.W.W., Fields, M.W., Alm, E.J., Hazen, T.C., Adams, P.D., Arkin, A.P. and Zhou, J. 2018. Microbial Functional Gene Diversity Predicts Groundwater Contamination and Ecosystem Functioning. mBio 9(1).

Hyatt, D., Chen, G.L., Locascio, P.F., Land, M.L., Larimer, F.W. and Hauser, L.J. 2010. Prodigal: prokaryotic gene recognition and translation initiation site identification. BMC Bioinformatics 11, 119.

Kanehisa, M. and Sato, Y. 2020. KEGG Mapper for inferring cellular functions from protein sequences. Protein Sci 29(1), 28–35.

Karthikeyan, O.P., Smith, T.J., Dandare, S.U., Parwin, K.S., Singh, H., Loh, H.X., Cunningham, M.R., Williams, P.N., Nichol, T., Subramanian, A., Ramasamy, K. and Kumaresan, D. 2021. Metal(loid) speciation and transformation by aerobic methanotrophs. Microbiome 9(1), 156.

King, A.J., Preheim, S.P., Bailey, K.L., Robeson, M.S., 2nd, Roy Chowdhury, T., Crable, B.R., Hurt, R.A., Jr., Mehlhorn, T., Lowe, K.A., Phelps, T.J., Palumbo, A.V., Brandt, C.C., Brown, S.D., Podar, M., Zhang, P., Lancaster, W.A., Poole, F., Watson, D.B., M, W.F., Chandonia, J.M., Alm, E.J., Zhou, J., Adams, M.W., Hazen, T.C., Arkin, A.P. and Elias, D.A. 2017. Temporal Dynamics of In-Field Bioreactor Populations Reflect the Groundwater System and Respond Predictably to Perturbation. Environ Sci Technol 51(5), 2879–2889.

Kodama, Y. and Watanabe, K. 2004. Sulfuricurvum kujiense gen. nov., sp. nov., a facultatively anaerobic, chemolithoautotrophic, sulfur-oxidizing bacterium isolated from an underground crude-oil storage cavity. Int J Syst Evol Microbiol 54(Pt 6), 2297–2300.

Kojima, H., Shinohara, A. and Fukui, M. 2015. Sulfurifustis variabilis gen. nov., sp. nov., a sulfur oxidizer isolated from a lake, and proposal of Acidiferrobacteraceae fam. nov. and Acidiferrobacterales ord. nov. Int J Syst Evol Microbiol 65(10), 3709–3713.

Liang, T., Ding, H., Wang, G., Kang, J., Pang, H. and Lv, J. 2016. Sulfur decreases cadmium translocation and enhances cadmium tolerance by promoting sulfur assimilation and glutathione metabolism in Brassica chinensis L. Ecotoxicol Environ Saf 124, 129–137.

Louca, S., Parfrey, L.W. and Doebeli, M. 2016. Decoupling function and taxonomy in the global ocean microbiome. Science 353(6305), 1272–1277.

Lozupone, C., Lladser, M.E., Knights, D., Stombaugh, J. and Knight, R. 2011. UniFrac: an effective distance metric for microbial community comparison. ISME J 5(2), 169–172.

Ma, X., Zhang, Q., Zheng, M., Gao, Y., Yuan, T., Hale, L., Van Nostrand, J.D., Zhou, J., Wan, S. and Yang, Y. 2019. Microbial functional traits are sensitive indicators of mild disturbance by lamb grazing. ISME J 13(5), 1370–1373.

Maestre, F.T., Delgado-Baquerizo, M., Jeffries, T.C., Eldridge, D.J., Ochoa, V., Gozalo, B., Quero, J.L., Garcia-Gomez, M., Gallardo, A., Ulrich, W., Bowker, M.A., Arredondo, T., Barraza-Zepeda, C., Bran, D., Florentino, A., Gaitan, J., Gutierrez, J.R., Huber-Sannwald, E., Jankju, M., Mau, R.L., Miriti, M., Naseri, K., Ospina, A., Stavi, I., Wang, D., Woods, N.N., Yuan, X., Zaady, E. and Singh, B.K. 2015. Increasing aridity reduces soil microbial diversity and abundance in global drylands. Proc Natl Acad Sci U S A 112(51), 15684–15689.

Nielsen, U. N., Ayres, E., Wall, D. H., & Bardgett, R. D. 2011. Soil biodiversity and carbon cycling: a review and synthesis of studies examining diversity–function relationships. Eur J Soil Sci, 62(1), 105–116.

Ning, D., Wang, Y., Fan, Y., Wang, J., Van Nostrand, J.D., Wu, L., Zhang, P., Curtis, D.J., Tian, R., Lui, L., Hazen, T.C., Alm, E.J., Fields, M.W., Poole, F., Adams, M.W.W., Chakraborty, R., Stahl, D.A., Adams, P.D., Arkin, A.P., He, Z. and Zhou, J. 2024. Environmental stress mediates groundwater microbial community assembly. Nat Microbiol.

Nunes, I., Jacquiod, S., Brejnrod, A., Holm, P.E., Johansen, A., Brandt, K.K., Prieme, A. and Sorensen, S.J. 2016. Coping with copper: legacy effect of copper on potential activity of soil bacteria following a century of exposure. FEMS Microbiol Ecol 92(11).

Nurk, S., Meleshko, D., Korobeynikov, A. and Pevzner, P.A. 2017. metaSPAdes: a new versatile metagenomic assembler. Genome Res 27(5), 824–834.

Oksanen, J., Blanchet, F. G., Friendly, M., Kindt, R., Legendre, P., McGlinn, D., Minchin, P. R., O’Hara, R. B., Simpson, G. L., Solymos, P., Henry, M., Stevens, H., Szoecs, E., & Wagner, H. 2020. Vegan: community ecology package. R package version 2.5–7. Available at https://CRAN.Rproject.org/package=vegan.

Poghosyan, L., Koch, H., Lavy, A., Frank, J., van Kessel, M., Jetten, M.S.M., Banfield, J.F. and Lucker, S. 2019. Metagenomic recovery of two distinct comammox Nitrospira from the terrestrial subsurface. Environ Microbiol 21(10), 3627–3637.

Power, J.F., Carere, C.R., Lee, C.K., Wakerley, G.L.J., Evans, D.W., Button, M., White, D., Climo, M.D., Hinze, A.M., Morgan, X.C., McDonald, I.R., Cary, S.C. and Stott, M.B. 2018. Microbial biogeography of 925 geothermal springs in New Zealand. Nat Commun 9(1), 2876.

Pradhan, S.K., Singh, N.R., Kumar U., Mishra, S.R., Perumal, R.C., Benny, J., Thatoi, H. 2020. Illumina MiSeq based assessment of bacterial community structure and diversity along the heavy metal concentration gradient in Sukinda chromite mine area soils. India. Ecological Genetics and Genomics, 15, 100054.

Prakash, O., Green, S.J., Singh, P., Jasrotia, P. and Kostka, J.E. 2021. Stress-related ecophysiology of members of the genus Rhodanobacter isolated from a mixed waste contaminated subsurface. Frontiers of Environmental Science & Engineering 15(2).

Price, M.N., Dehal, P.S. and Arkin, A.P. 2009. FastTree: computing large minimum evolution trees with profiles instead of a distance matrix. Mol Biol Evol 26(7), 1641–1650.

Price, M.N., Dehal, P.S. and Arkin, A.P. 2010. FastTree 2--approximately maximum-likelihood trees for large alignments. PLoS One 5(3), e9490.

Rocca, J.D., Simonin, M., Blaszczak, J.R., Ernakovich, J.G., Gibbons, S.M., Midani, F.S. and Washburne, A.D. 2018. The Microbiome Stress Project: Toward a Global Meta-Analysis of Environmental Stressors and Their Effects on Microbial Communities. Front Microbiol 9, 3272.

Ruhl, I.A., Grasby, S.E., Haupt, E.S. and Dunfield, P.F. 2018. Analysis of microbial communities in natural halite springs reveals a domain-dependent relationship of species diversity to osmotic stress. Environ Microbiol Rep 10(6), 695–703.

Shi, Z. J., Xiao, N., Ning, D., Tian, R., Zhang, P., Curtis, D., Van Nostrand, J.D., Wu, L., Hazen, T.C., Rocha, A.M., He, Z., Arkin, A.P., Firestone, M.K., Zhou, J. 2022. EcoFun-MAP: An Ecological Function Oriented Metagenomic Analysis Pipeline. 10.1101/2022.04.05.481366.

Smith, M.B., Rocha, A.M., Smillie, C.S., Olesen, S.W., Paradis, C., Wu, L., Campbell, J.H., Fortney, J.L., Mehlhorn, T.L., Lowe, K.A., Earles, J.E., Phillips, J., Techtmann, S.M., Joyner, D.C., Elias, D.A., Bailey, K.L., Hurt, R.A., Jr., Preheim, S.P., Sanders, M.C., Yang, J., Mueller, M.A., Brooks, S., Watson, D.B., Zhang, P., He, Z., Dubinsky, E.A., Adams, P.D., Arkin, A.P., Fields, M.W., Zhou, J., Alm, E.J. and Hazen, T.C. 2015. Natural bacterial communities serve as quantitative geochemical biosensors. mBio 6(3), e00326–00315.

Song, H.K., Shi, Y., Yang, T., Chu, H., He, J.S., Kim, H., Jablonski, P. and Adams, J.M. 2019. Environmental filtering of bacterial functional diversity along an aridity gradient. Sci Rep 9(1), 866.

Song, L., Pan, Z., Dai, Y., Chen, L., Zhang, L., Liao, Q., Yu, X., Guo, H. and Zhou, G. 2020. Characterization and comparison of the bacterial communities of rhizosphere and bulk soils from cadmium-polluted wheat fields. PeerJ 8, e10302.

Talbot, J.M., Bruns, T.D., Taylor, J.W., Smith, D.P., Branco, S., Glassman, S.I., Erlandson, S., Vilgalys, R., Liao, H.L., Smith, M.E. and Peay, K.G. 2014. Endemism and functional convergence across the North American soil mycobiome. Proc Natl Acad Sci U S A 111(17), 6341–6346.

Thevenot, E.A., Roux, A., Xu, Y., Ezan, E. and Junot, C. 2015. Analysis of the Human Adult Urinary Metabolome Variations with Age, Body Mass Index, and Gender by Implementing a Comprehensive Workflow for Univariate and OPLS Statistical Analyses. J Proteome Res 14(8), 3322–3335.

Versantvoort, W., Guerrero-Cruz, S., Speth, D.R., Frank, J., Gambelli, L., Cremers, G., van Alen, T., Jetten, M.S.M., Kartal, B., Op den Camp, H.J.M. and Reimann, J. 2018. Comparative Genomics of Candidatus Methylomirabilis Species and Description of Ca. Methylomirabilis Lanthanidiphila. Front Microbiol 9, 1672.

Wang, Y., Feng, H., Wang, R., Zhou, L., Li, N., He, Y., Yang, X., Lai, J., Chen, K. and Zhu, W. 2023. Non-targeted metabolomics and 16s rDNA reveal the impact of uranium stress on rhizosphere and non-rhizosphere soil of ryegrass. J Environ Radioact 258, 107090.

Warton, D.I., Wright, S.T. and Wang, Y. 2012. Distance-based multivariate analyses confound location and dispersion effects. Methods in Ecology and Evolution 3(1), 89–101.

Watanabe, T., Miura, A., Iwata, T., Kojima, H. and Fukui, M. 2017. Dominance of Sulfuritalea species in nitrate-depleted water of a stratified freshwater lake and arsenate respiration ability within the genus. Environ Microbiol Rep 9(5), 522–527.

Wu, L., Wen, C., Qin, Y., Yin, H., Tu, Q., Van Nostrand, J.D., Yuan, T., Yuan, M., Deng, Y. and Zhou, J. 2015. Phasing amplicon sequencing on Illumina Miseq for robust environmental microbial community analysis. BMC Microbiol 15, 125.

Whittaker, R. H. 1972. Evolution and measurement of species diversity. Taxon 21(2-3), 213–251.

Xu, M., Wu, W.M., Wu, L., He, Z., Van Nostrand, J.D., Deng, Y., Luo, J., Carley, J., Ginder-Vogel, M., Gentry, T.J., Gu, B., Watson, D., Jardine, P.M., Marsh, T.L., Tiedje, J.M., Hazen, T., Criddle, C.S. and Zhou, J. 2010. Responses of microbial community functional structures to pilot-scale uranium in situ bioremediation. ISME J 4(8), 1060–1070.

Zaneveld, J.R., McMinds, R. and Vega Thurber, R. 2017. Stress and stability: applying the Anna Karenina principle to animal microbiomes. Nat Microbiol 2, 17121.

Zhang, P., Wu, W.M., Van Nostrand, J.D., Deng, Y., He, Z., Gihring, T., Zhang, G., Schadt, C.W., Watson, D., Jardine, P., Criddle, C.S., Brooks, S., Marsh, T.L., Tiedje, J.M., Arkin, A.P. and Zhou, J. 2015. Dynamic Succession of Groundwater Functional Microbial Communities in Response to Emulsified Vegetable Oil Amendment during Sustained In Situ U(VI) Reduction. Appl Environ Microbiol 81(12), 4164–4172.

Zhao, S., Ye, Z. and Stanton, R. 2020. Misuse of RPKM or TPM normalization when comparing across samples and sequencing protocols. RNA 26(8), 903–909.

Zhou, J., Deng, Y., Zhang, P., Xue, K., Liang, Y., Van Nostrand, J.D., Yang, Y., He, Z., Wu, L., Stahl, D.A., Hazen, T.C., Tiedje, J.M. and Arkin, A.P. 2014. Stochasticity, succession, and environmental perturbations in a fluidic ecosystem. Proc Natl Acad Sci U S A 111(9), E836–845.

Zhou, J. and Ning, D. 2017. Stochastic Community Assembly: Does It Matter in Microbial Ecology? Microbiol Mol Biol Rev 81(4).

