## Supplemental Figure 1 to Figure 4 and supplemental table 1 to 2 for "Modest functional diversity decline and pronounced composition shifts of microbial communities in a uranium-contaminated aquifer"

Table S1. Aquifer environmental variables in in uncontaminated wells (UC), low-contaminated wells (LC), mid-contaminated wells (MC), and high-contaminated wells (HC).

|  | UC | LC | MC | HC |
| --- | --- | --- | --- | --- |
| Conductivity (μS/cm) | 242.53±175.75^b^ | 452±183.92^b^ | 1000.8±609.1^b^ | 9830.33±3316.99^a^ |
| pH | 6.43±0.94^b^ | 6.62±0.08^b^ | 6.58±0.1^b^ | 4.06±1^a^ |
| Redox (mV) | 42.33±151.79^ab^ | 474.33±287.25^ab^ | 32.67±78.01^a^ | 332.67±129.36^b^ |
| DO (mg/L) | 1.87±1.65^a^ | 2.06±2.96^a^ | 0.27±0.14^a^ | 0.54±0.55^a^ |
| DIC (mg/L) | 34.21±17.03^a^ | 54.29±28.13^a^ | 74.96±20.17^a^ | 60.81±49.77^a^ |
| DOC (mg/L) | 15.24±25.38^ab^ | 0.86±0.44^a^ | 1.95±0.37^b^ | 20.56±23.65^ab^ |
| O_2_ (mM) | 0.25±0.12^a^ | 0.5±0.25^a^ | 0.23±0.09^a^ | 0.17±0.06^a^ |
| CH_4_ (mM) | 0.23±0.23^a^ | 0.02±0.02^a^ | 0.02±0.006^a^ | 0 |
| CO_2_ (mM) | 12.05±18.25^ab^ | 3.49±1.41^b^ | 4.4±1.57^b^ | 15.06±5.84^a^ |
| N_2_ (mM) | 1.42±0.45^ab^ | 2.48±0.95^a^ | 1.11±0.39^ab^ | 0.69±0.24^b^ |
| N_2_O (mM) | 0.11±0.11^a^ | 0 | 0.1±0.1^a^ | 0.88±0.27^b^ |
| S^2-^ (mg/L) | 0.026±0.026^a^ | 0.002±0.001^a^ | 0.001±0.001^a^ | 0.002±0.002^a^ |
| Cl^-^ (mg/L) | 1.63±0.49^c^ | 12.35±15.65^bc^ | 21.04±4.72^b^ | 238.36±83.1^a^ |
| NO_3_^-^ (mg/L) | 1.36±2^b^ | 1.16±1.87^b^ | 321.03±434.46^ab^ | 5422.55±3285.46^a^ |
| PO_4_^-^ (mg/L) | ND | 0.08±0.13 | ND | ND |
| SO_4_^2-^ (mg/L) | 7.16±0.91^a^ | 15±5.16^a^ | 49.48±36.62^a^ | 799.51±1101.18^a^ |
| NO_3__sup (mM) | ND | 0.02±0.04^a^ | 3.25±5.56^a^ | 38.65±33.63^a^ |
| NO_2__sup (µM) | ND | ND | 19.51±24.26^a^ | 13.61±4.49^a^ |
| Ferrous (mg/L) | 0.18±0.28^a^ | 0.33±0.45^a^ | 0.56±0.79^a^ | 1.08±1.07^a^ |
| K (mg/L) | 1.3±0.29^a^ | 3.09±1.59^ab^ | 4.8±1.38^b^ | 111.38±95.81^ab^ |
| Na (mg/L) | ND | ND | 16.35±14.93^a^ | 508.44±315^a^ |
| Ca (mg/L) | 41.29±33.12^a^ | 69.19±26.08^a^ | 704.8±995.72^a^ | 5463.17±6075.62^a^ |
| Mg (mg/L) | ND | ND | 9.96±17.25^a^ | 117.05±81.56^a^ |
| Al (mg/L) | 0.01±0.02^a^ | 0.6±1.01^a^ | 1.15±1.99^a^ | 76.27±61.1^a^ |
| Ba (mg/L) | 0.07±0.02^a^ | 0.1±0.01^a^ | 0.33±0.26^a^ | 0.82±1.09^a^ |
| Ag (mg/L) | 0.005±0.004^b^ | 0.01±0.0007^ab^ | 0.007±0.006^ab^ | 0.018±0.006^a^ |
| As (mg/L) | ND | 0.0006±0.001^a^ | ND | 0.003±0.005^a^ |
| Be (mg/L) | ND | 0.006±0.01^a^ | ND | 0.06±0.02^b^ |
| Bi (mg/L) | ND | 0.001±0.005 | ND | ND |
| Cd (mg/L) | ND | 0.002±0.001^a^ | 0.003±0.006^a^ | 0.142±0.03^b^ |
| Co (mg/L) | ND | 0.01±0.02^a^ | 0.02±0.03^a^ | 0.62±0.55^a^ |
| Cr (mg/L) | ND | 0.004±0.002^a^ | 0.002±0.003^a^ | 0.13±0.22^a^ |
| Cs (mg/L) | ND | 0.003±0.005^a^ | ND | 0.011±0.01^a^ |
| Cu (mg/L) | ND | 0.014±0.002^a^ | ND | 0.32±0.42^a^ |
| Zn (mg/L) | 0.06±0.01^a^ | 0.06±0.02^a^ | 0.05±0.02^a^ | 0.73±0.47^a^ |
| Fe (mg/L) | 0.02±0.04^a^ | 0.76±0.41^a^ | 0.64±1.12^a^ | 1.08±1.84^a^ |
| Ga (mg/L) | ND | 0.005±0.004^a^ | 0.011±0.01^a^ | 0.03±0.02^a^ |
| Mn (mg/L) | 0.07±0.11^a^ | 1.58±2.48^a^ | 3.38±5.21^a^ | 95±54.18^b^ |
| Li (mg/L) | ND | 0.02±0.01^a^ | 0.016±0.03^a^ | 0.75±1.04^a^ |
| Ni (mg/L) | ND | 0.051±0.09^a^ | 0.081±0.14^a^ | 4.52±3.15^a^ |
| Pb (mg/L) | ND | 0.0008±0.0007^a^ | ND | 0.013±0.016^a^ |
| Se (mg/L) | ND | ND | ND | 0.012±0.012 |
| Sr (mg/L) | 0.15±0.04^a^ | 0.10±0.11^a^ | 0.52±0.23^a^ | 1.7±1.52^a^ |
| U (mg/L) | 0.07±0.12^a^ | 0.004±0.004^b^ | 0.75±0.7^ab^ | 9.79±6.47^ab^ |
| Mo^95^_sup (µM) | 0.002±0.003^a^ | 0.001±0.0009^a^ | 0.01±0.02^a^ | 0.004±0.007^a^ |

^a^Letters represent the difference between groups. The difference between groups was determined by ANOVA. Significant difference level: *p* < 0.05. ND: Not detected; sup: concentration in the supernatant of each sample.

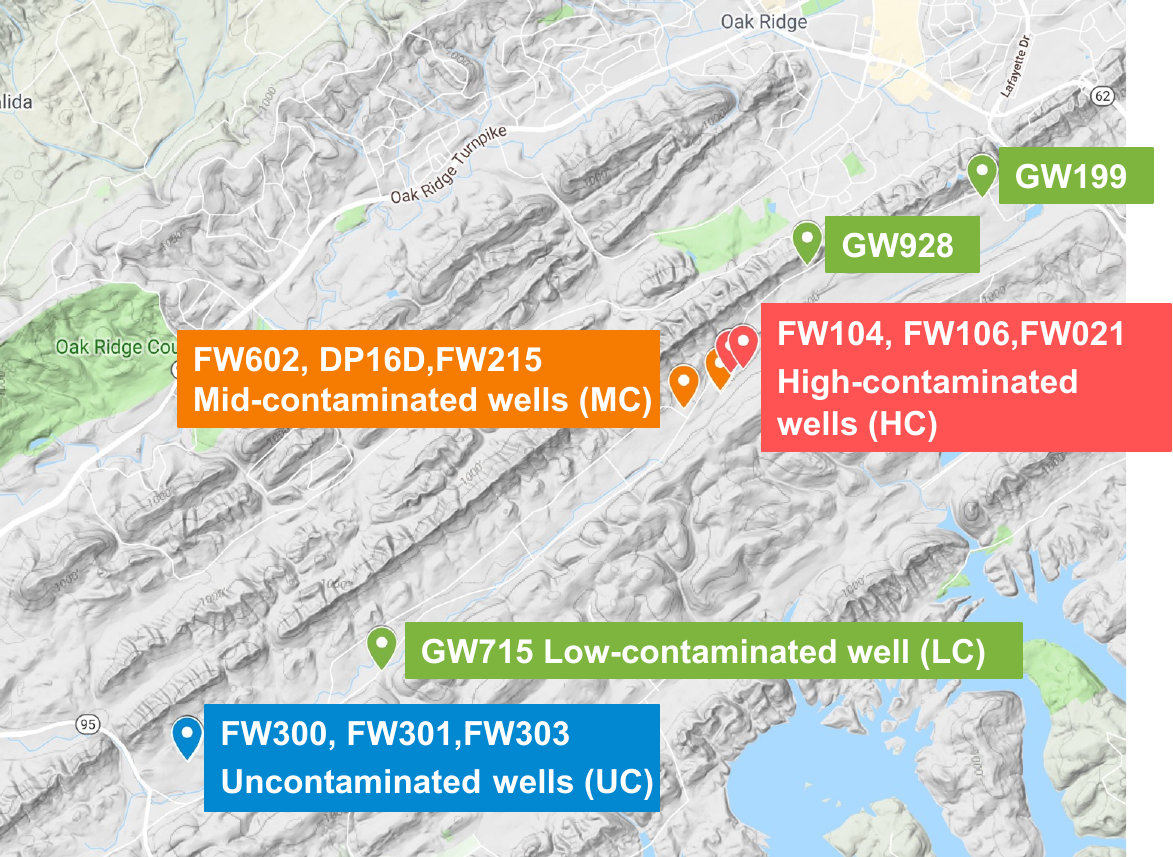

Fig. S1. Geographical location of the study sites. Aquifer samples consist of uncontaminated wells (UC) FW300, FW301, FW303, and FW305; low-contaminated wells (LC) GW199, GW715, and GW928; mid-contaminated wells (MC) FW215, FW602, and DP16D; high-contaminated wells (HC) FW104, FW106, and FW021.

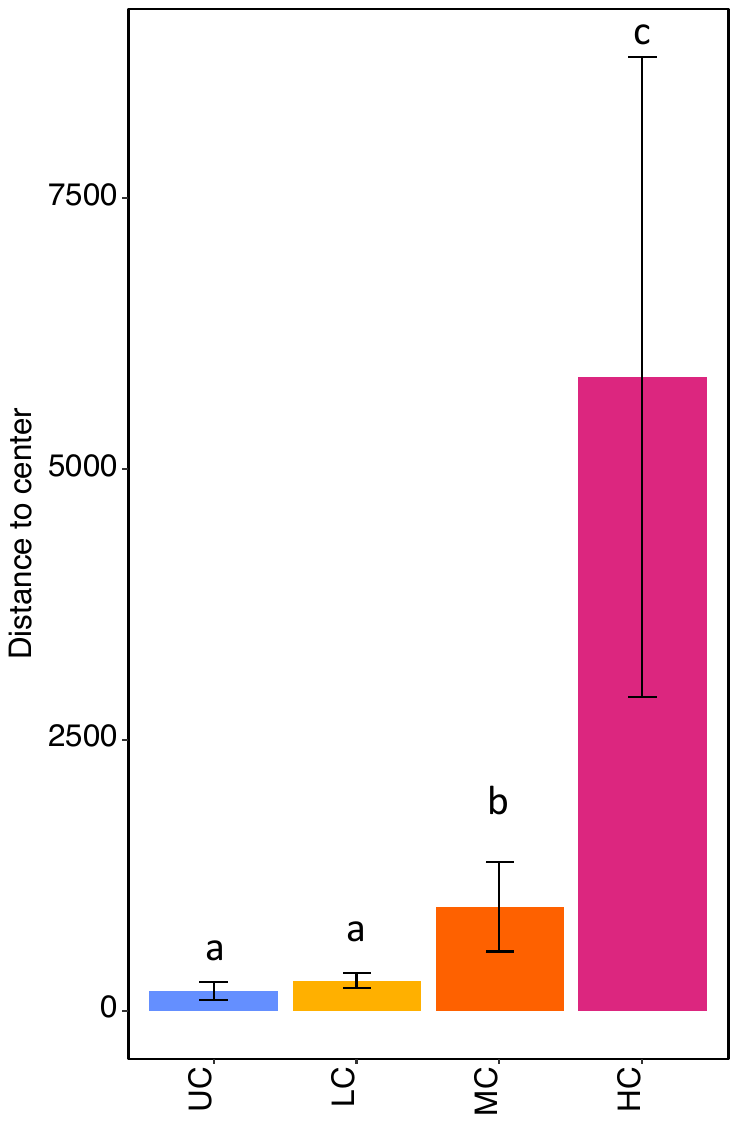

Fig. S2. Dispersion analysis of environmental variables based on Euclidean distance. Letters represent the difference between groups which was determined by ANOVA (significant difference level: *p* < 0.05).

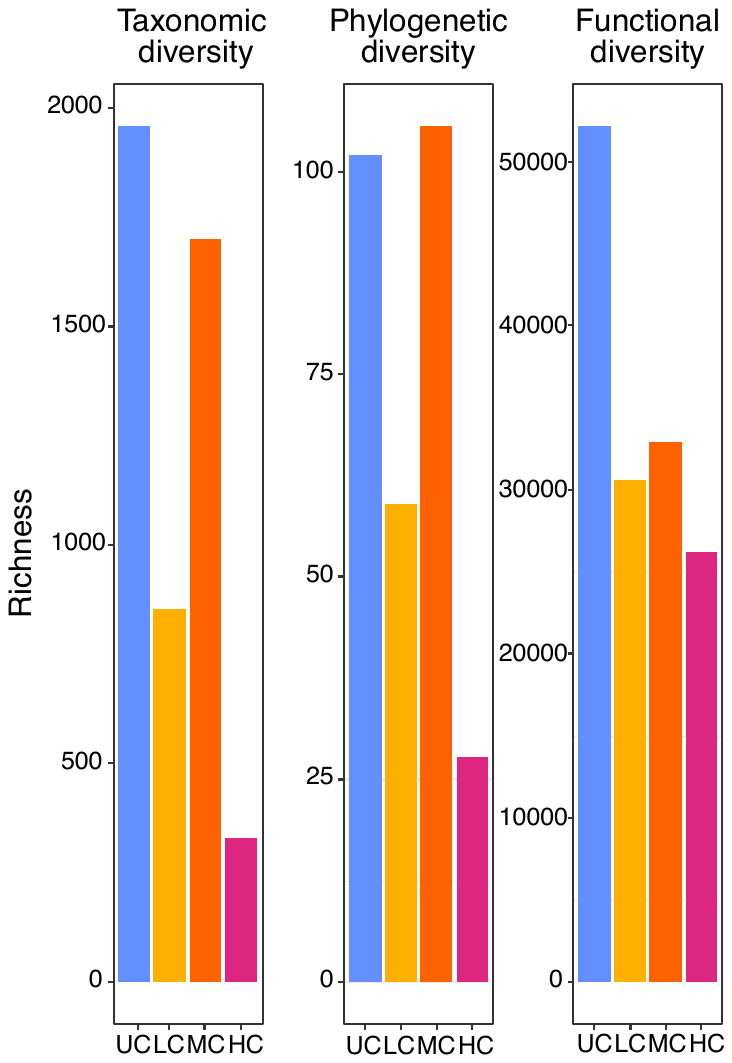

Fig. S3. Microbial γ-diversity of 12 groundwater samples calculated by richness.

Table S2. Taxonomic composition of microbial communities at phylum (A) and genus (B) level under different treatments.

| **A**.Category at Phylum level | UC | LC | MC | HC |
| --- | --- | --- | --- | --- |
|  | Avergae relative abundance ± standard deviation (%) | | | |
| Gammaproteobacteria | 42.86±19.18^a^ | 27.21±15.73^a^ | 23.45±1.90^a^ | 73.06±22.44^a^ |
| Alphaproteobacteria | 12.97±7.44^b^ | 18.39±11.88^a^ | 3.36±0.41^cd^ | 1.25±1.89^b^ |
| Verrucomicrobiota | 4.76±3.14^b^ | 3.97±2.44^a^ | 16.29±4.36^ab^ | 4.54±7.74^b^ |
| Bacteroidota | 3.22±3.10^b^ | 11.72±6.57^a^ | 5.15±2.62^bc^ | 1.64±1.37^b^ |
| Acidobacteriota | 2.20±1.08^b^ | 0.83±0.56^b^ | 6.33±5.11^bc^ | 5.47±7.40^b^ |
| WPS-2 | 0.22±0.23^b^ | 0.07±0.08^b^ | 0.54±0.74^c^ | 12.45±21.56^b^ |
| Nitrospirota | 2.54±2.75^b^ | 6.72±6.98^a^ | 2.57±1.90^c^ | 0.06±0.05^b^ |
| Patescibacteria | 3.43±2.01^b^ | 0.41±0.30^b^ | 7.02±4.53^bc^ | 0.31±0.51^b^ |
| Bdellovibrionota | 8.39±10.41^b^ | 0.84±1.22^b^ | 0.96±1.01^c^ | 0.00±0.00^b^ |
| Myxococcota | 1.13±0.97^b^ | 1.92±2.55^b^ | 5.97±2.90^bc^ | 0.02±0.02^b^ |
| Methylomirabilota | 0.70±0.89^b^ | 0.22±0.30^b^ | 7.04±7.13^bc^ | 0.01±0.01^b^ |
| Elusimicrobiota | 4.06±2.44^b^ | 0.81±1.10^b^ | 1.58±1.37^c^ | 0.01±0.01^b^ |
| Campilobacterota | 6.06±10.32^b^ | 0.00±0.00^b^ | 0.02±0.01^c^ | 0.00±0.00^b^ |
| Chloroflexi | 0.46±0.41^b^ | 0.70±0.98^b^ | 4.01±5.90^cd^ | 0.11±0.15^b^ |
| DTB120 | 0.12±0.18^b^ | 0.04±0.06^b^ | 3.73±5.58^cd^ | 0.12±0.21^b^ |
| Actinobacteriota | 0.60±0.51^b^ | 2.04±1.81^a^ | 0.72±0.42^c^ | 0.12±0.08^b^ |
| Spirochaetota | 0.09±0.06^b^ | 0.01±0.02^b^ | 2.66±2.62^c^ | 0.14±0.24^b^ |
| Crenarchaeota | 0.20±0.05^b^ | 22.93±30.70^a^ | 0.17±0.18^c^ | 0.00±0.00^b^ |
| Others | 6.00±2.45 ^b^ | 1.16±0.57^b^ | 8.45±3.46^bc^ | 0.69±0.78^b^ |

| **B**.Category at Genus level | UC | LC | MC | HC |
| --- | --- | --- | --- | --- |
|  | Avergae relative abundance ± standard deviation (%) | | | |
| Rhodanobacter | 0.01±0.02^c^ | 0.02±0.03^b^ | 0.86±0.89^c^ | 37.50±35.73^a^ |
| Nitrosarchaeum | 0.06±0.03^c^ | 19.18±33.14^ab^ | 0.14±0.20^c^ | 0.00±0.00^b^ |
| Candidatus_Omnitrophus | 1.14±1.81^c^ | 3.14±2.60^b^ | 10.05±8.20^bc^ | 3.93±6.72^ab^ |
| WPS-2 | 0.22±0.23^c^ | 0.07±0.08^b^ | 0.54±0.74^c^ | 12.45±21.56^ab^ |
| Sulfuritalea | 1.72±0.99^c^ | 9.85±8.40^ab^ | 0.07±0.12^c^ | 0.00±0.00^b^ |
| Sulfurifustis | 0.44±0.56^c^ | 0.10±0.09^b^ | 1.56±1.57^bc^ | 9.51±16.46^ab^ |
| Chujaibacter | 0.00±0.00^c^ | 0.00±0.00^b^ | 1.01±1.09^c^ | 7.52±13.03^ab^ |
| Reyranella | 0.84±1.08^c^ | 6.25±7.19^b^ | 0.13±0.13^c^ | 0.48±0.82^b^ |
| GOUTA6 | 0.90±1.19^c^ | 0.18±0.32^b^ | 5.47±1.33^bc^ | 0.40±0.70^b^ |
| Candidatus_Methylomirabilis | 0.00±0.00^c^ | 0.03±0.04^b^ | 6.79±7.08^bc^ | 0.01±0.01^b^ |
| Nitrospira | 1.24±1.19^c^ | 5.35±5.37^b^ | 0.23±0.31^c^ | 0.00±0.00^b^ |
| 0319-6G20 | 5.93±7.67^bc^ | 0.22±0.35^b^ | 0.58±0.49^c^ | 0.00±0.00^b^ |
| bacteriap25 | 0.55±0.60^c^ | 0.55±0.86^b^ | 5.50±2.94^bc^ | 0.00±0.00^b^ |
| Brevundimonas | 1.51±1.76^c^ | 4.79±5.84^b^ | 0.04±0.07^c^ | 0.00±0.00^b^ |
| Sulfuricurvum | 5.90±10.04^bc^ | 0.01±0.01^b^ | 0.00±0.01^c^ | 0.00±0.00^b^ |
| Omnitrophales | 0.96±0.60^c^ | 0.28±0.41^b^ | 4.34±5.83^bc^ | 0.00±0.00^b^ |
| Sediminibacterium | 1.49±2.21^c^ | 3.45±2.04^b^ | 0.00±0.01^c^ | 0.22±0.38^b^ |
| Others | 46.78±4.31^a^ | 34.94±8.97^a^ | 42.62±7.12^a^ | 14.90±14.43^ab^ |
| unclassified | 18.24±12.41^b^ | 7.18±4.16^b^ | 8.34±2.45^bc^ | 11.24±5.96^ab^ |
| uncultured | 12.08±8.58^bc^ | 4.40±2.60^b^ | 11.71±3.03^b^ | 1.83±1.50^b^ |

^a^Letters represent the difference between groups. The difference between groups was determined by ANOVA. Significant difference level: *p* < 0.05.

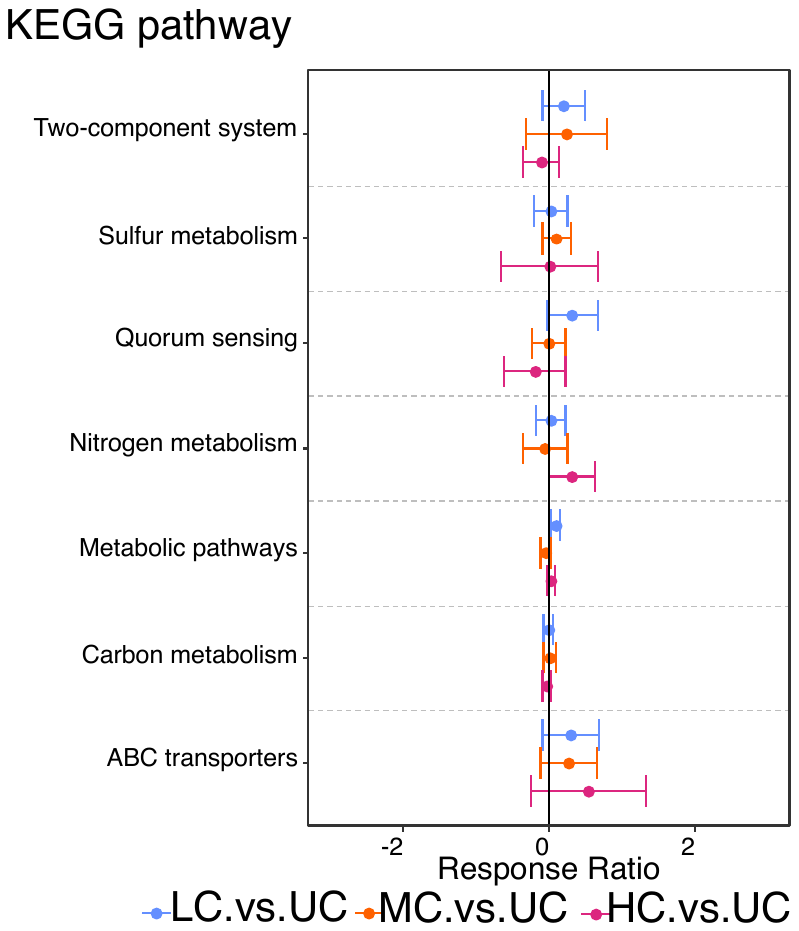

Fig. S4. Differences in assembly-based shotgun metagenomic data between contaminated wells and uncontaminated wells, reflected by response ratios of annotated KEGG pathway.
